# Rebuilding microbiome diversity theory on the closed simplex

**DOI:** 10.64898/2026.09.02.748976

**Authors:** Yiqian Zhang, Zihan Zhu

## Abstract

Ecological diversity theory links diversity within local communities to diversity of higher-level ensembles, but this scale structure is largely absent from microbiome analysis. Alpha diversity is usually treated as a within-sample summary, whereas “beta diversity” often denotes pairwise dissimilarity and gamma diversity is rarely explicit. We restore the local-regional architecture for environmental, host-associated and longitudinal microbiomes and distinguish regional beta diversity from pairwise dissimilarity and predictor-associated compositional variation. To quantify these objects for sparse compositions, we introduce Hellinger-Riemann intrinsic coordinates (HRIC), a one-to-one, bounded normal-coordinate representation of the closed simplex that retains exact zeros. The same coordinates yield Simplex Hellinger alpha and gamma diversity, additive regional beta diversity, taxon contributions, pairwise dissimilarity and model-explained dispersion. Simulations established the correspondence between HRIC dispersion and the between-condition component of PERMANOVA. Across Arctic and North Atlantic communities, local diversity relative to each regional benchmark covaried similarly with vertical environmental gradients despite partly different taxon-level associations. In a randomized autologous faecal microbiota transplantation trial, recipients returned earlier towards their personal pre-transplant compositions, whereas the alpha-diversity difference was smaller and less precise. Explicit local and regional referents therefore connect diversity partitioning with compositional analysis across microbial systems.

## 1 Introduction

Ecological diversity theory describes how diversity is organized within local communities and across higher-level ensembles. In environmental microbiology, diversity analyses relate community organization across habitats and physicochemical gradients to ecosystem processes [1, 2]. In host-associated microbiome research, they characterize body-site specialization, interpersonal variation, dysbiosis and recovery after perturbation [3, 4]. Yet contemporary microbiome studies typically treat alpha diversity as a within-sample summary and “beta diversity” as between-sample dissimilarity, while gamma diversity and the regional ensemble that gives local–regional beta diversity its ecological meaning are rarely explicit [3, 5].

This contraction accompanied the translation of ecological terminology into sequencing-based workflows. UniFrac made phylogenetic pairwise dissimilarity available for community comparison [6]; QIIME integrated alpha diversity with dissimilarity-based analysis [5]; and the Human Microbiome Project carried this template to population-scale data, contrasting alpha diversity with Bray–Curtis “beta diversity” among subjects [3]. In Whittaker’s framework, however, alpha diversity describes diversity within local communities, gamma diversity describes diversity of the pooled regional composition, and regional beta diversity describes differentiation between these scales [7]. Hill and Jost later formalized this architecture in effective-number terms [8, 9]. Alongside it, a related tradition developed from pair-wise resemblance and gradient ordination to compositional variance, multivariate dispersion and distance-based association [7, 10–12]. Regional beta diversity, pairwise dissimilarity and predictor-associated dispersion therefore answer distinct ecological questions even when derived from a common representation. Moreover, regional beta diversity is meaningful only after the local community and higher-level ensemble have been specified: local units may be water samples, host-specific microbiota or time-specific specimens, but their regional grouping requires a defensible basis for joint interpretation and pooling.

Translating this architecture into quantitative analysis requires alpha and gamma diversity, regional beta diversity, pairwise dissimilarity and predictor-associated dispersion to retain a common representation of composition. This is challenging because microbial profiles are sparse compositions: their components carry relative information, sum to one and often contain exact zeros on boundary faces of the simplex. Hill numbers provide a coherent effective-number scale for alpha, regional beta and gamma diversity, but not sample coordinates for dissimilarities, centroids, ordinations or model-based sums of squares [8, 9]. Aitchison geometry supplies Euclidean log-ratio coordinates, but is defined on the strictly positive simplex, so observed zeros require replacement before transformation [13, 14]. A common representation for sparse microbial communities should instead operate directly on the closed simplex, preserve the supplied composition, keep boundary observations finite and support both diversity partitioning and multivariate analysis on a fixed feature dictionary.

We therefore extend the local–regional architecture to environmental, host-associated and longitudinal microbial communities and distinguish regional beta diversity from pairwise dissimilarity and predictor-associated dispersion. We introduce Hellinger–Riemann intrinsic coordinates (HRIC), a zero-compatible, one-to-one map from the closed simplex to a bounded subset of Euclidean contrast space. The same coordinates yield Simplex Hellinger alpha and gamma diversity, additive regional beta diversity, taxon contributions, pairwise dissimilarity and predictor-associated dispersion. We establish the framework mathematically, evaluate its operating characteristics in simulations and apply it to ocean environmental gradients and microbiota recovery after auto-FMT to distinguish scalar diversity organization from taxon-level and individualized compositional variation.

## 2 Results

### 2.1 A local–regional framework for microbial diversity and community differentiation

Classical local–regional diversity theory provides the basis for its extension to microbial ecology (Fig. 1a). Alpha, gamma and regional beta diversity describe diversity within local communities, diversity of the pooled regional composition and differentiation between these scales, respectively. Additive regional beta is gamma minus mean local alpha, whereas Hill–Jost multiplicative regional beta is the effective number of compositionally distinct communities [7–9, 15]. These architectures are not interchangeable, and both require specified local and regional units.

**Fig. 1.**
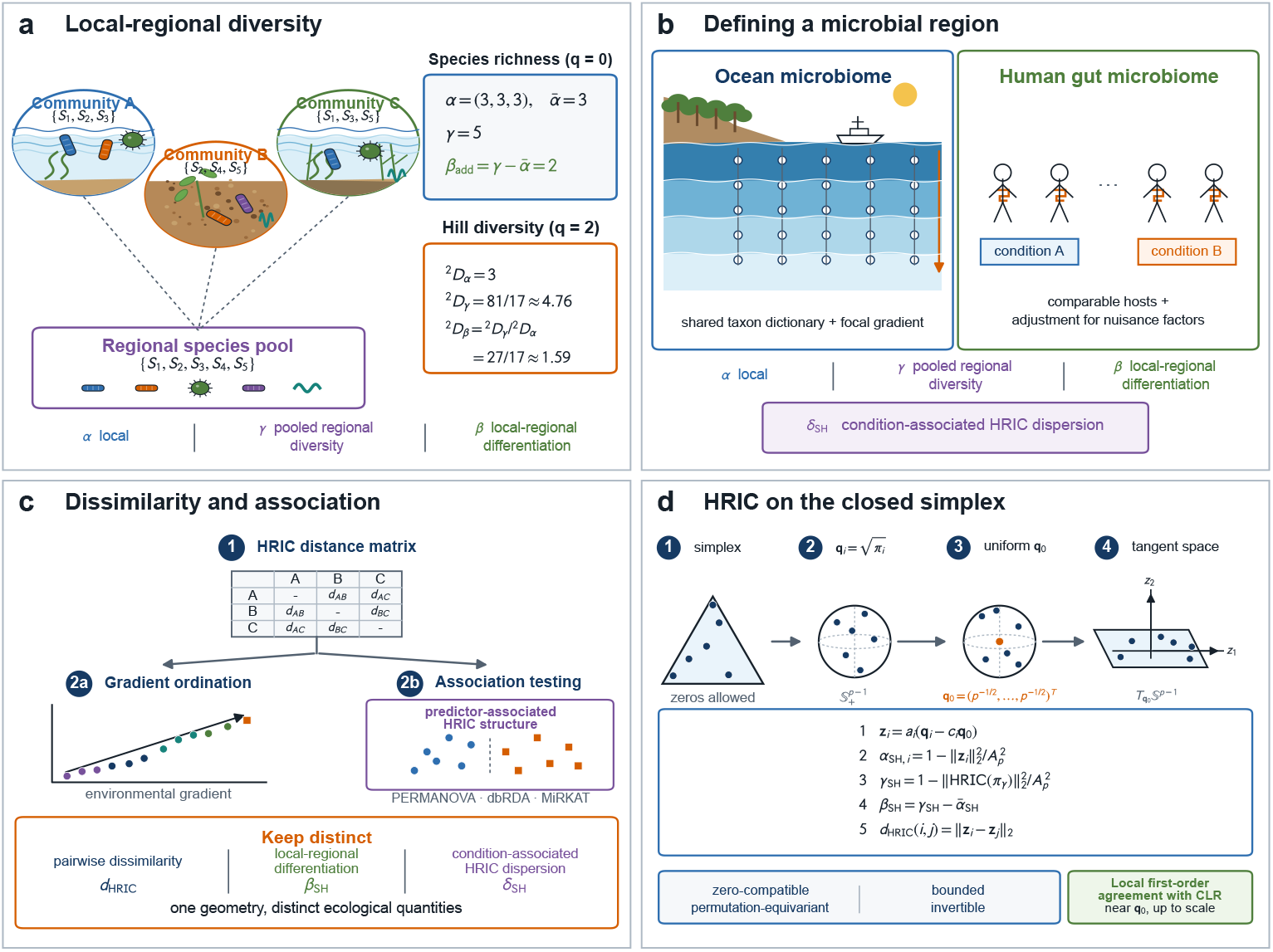
A common closed-simplex representation links local–regional microbial diversity and community differentiation. **a**, Local–regional diversity illustrated by a three-community richness example using additive and Whittaker multiplicative summaries, together with the corresponding order-2 Hill-number multiplicative partition. **b**, Operational definition of a microbial regional ensemble for ocean and host-associated microbiomes. Local compositions are pooled within a shared analytical and ecological context; condition-associated HRIC dispersion is distinguished from regional beta diversity. **c**, Development of dissimilarity-based community analysis from gradient ordination to contemporary association testing. Pairwise dissimilarity, regional beta diversity and condition-associated HRIC dispersion are defined within one geometry but answer different ecological questions. **d**, HRIC maps closed-simplex compositions through the square-root sphere into a Euclidean tangent space at the uniform composition, from which alpha and gamma diversity, regional beta diversity and pairwise dissimilarity are constructed.

For microbial ecology, we define a regional ensemble as an analytically prespecified collection of local communities for which joint interpretation and pooling are ecologically defensible (Fig. 1b). Local units may be water samples, host-specific microbiota or time-specific specimens. A region need not imply a specified dispersal process or demonstrably shared source pool, but quantitative comparison requires a common feature dictionary and an explicit weighting rule.

Regional beta diversity, pairwise dissimilarity and predictor-associated dispersion also differ in ecological referent (Fig. 1c). The first relates local to regional diversity, the second compares two compositions, and the third quantifies HRIC-coordinate dispersion explained by a specified predictor [7, 10, 12]. A common representation can connect these quantities without identifying them; their history is reviewed in Supplementary Note 1.

To quantify these objects jointly, we map each closed-simplex composition to Euclidean contrast space. For a composition ***π***_*i*_ = (*π*_*i*1_, …, *π*_*ip*_)^T^ of community *i* over *p* features, let 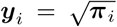 denote its element-wise square root on the non-negative orthant of the unit sphere, and let ***y***_0_ = *p*^*−*1*/*2^**1** denote the image of the uniform composition on that sphere [16]. The Hellinger–Riemann intrinsic coordinates are obtained by applying the spherical logarithmic map at ***y***_0_ to ***y***_*i*_,

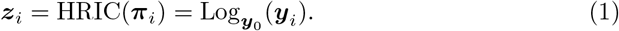

HRIC maps closed-simplex compositions, including exact-zero boundary points, one-to-one into a bounded subset of the tangent space at ***y***_0_. Its coordinates are zero-sum contrasts whose norm equals angular departure from uniformity. Further properties are established in Supplementary Note 2 and summarized in Supplementary Table 1 (Fig. 1d).

Within this representation, local diversity is measured by normalized radial departure from uniformity. Let 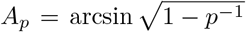 denote the maximum HRIC norm. Simplex Hellinger alpha diversity is

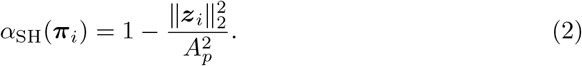

It ranges from 1 at uniformity to 0 at a simplex vertex and is a bounded monotonic reparameterization of order-1*/*2 Hill diversity (Supplementary Section S3.1). Its complement yields additive taxon contributions to community unevenness,

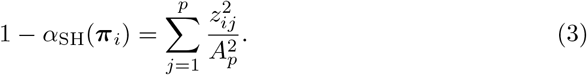

For a regional ensemble **Π** = (***π***_1_, …, ***π***_*n*_)^T^, let ***π***_*γ*_ be the normalized mean square-root composition under equal community weights. Simplex Hellinger gamma diversity is

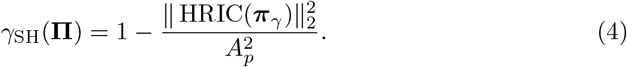

The corresponding additive regional beta diversity is

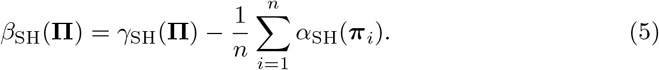

This non-negative difference is the increase in normalized diversity under pooling.

Pairwise HRIC dissimilarity is

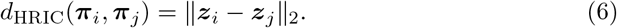

Unlike *β*_SH_, which partitions local and regional diversity, pairwise dissimilarity compares two coordinates and predictor-associated dispersion measures model-explained coordinate variation. The three quantities and their units of analysis are summarized in Supplementary Table 2.

Supplementary Notes 2 and 3 provide derivations and simulations. Under equal weights, condition-associated HRIC dispersion equals the PERMANOVA between-condition sum of squares divided by *n*. Simulations showed sensitivity primarily to compositional-mass redistribution and well-calibrated rejection under the null (Supplementary Section S3.11, Supplementary Fig. 1 and Supplementary Table 3).

### 2.2 Local–regional microbial diversity along environmental gradients in the Arctic Ocean and North Atlantic

The ocean water column provides an environmental setting in which microbial communities can be organized into local–regional ensembles. We reanalysed rRNA miTAG community profiles and matched environmental metadata from the Tara Oceans expedition [2, 17], defining 38 Arctic Ocean samples and 24 North Atlantic Ocean samples as two regional ensembles on the basis of their oceanographic context (Fig. 2 a,e). Individual water samples served as local communities. The two regions are geographically adjacent and connected by large-scale Atlantic–Arctic circulation, but differ in temperature, stratification, sea-ice influence and productivity regimes [18, 19]. We asked whether the two ensembles showed concordant associations between local–regional diversity and environmental gradients and whether the corresponding taxon-level associations were shared.

**Fig. 2.**
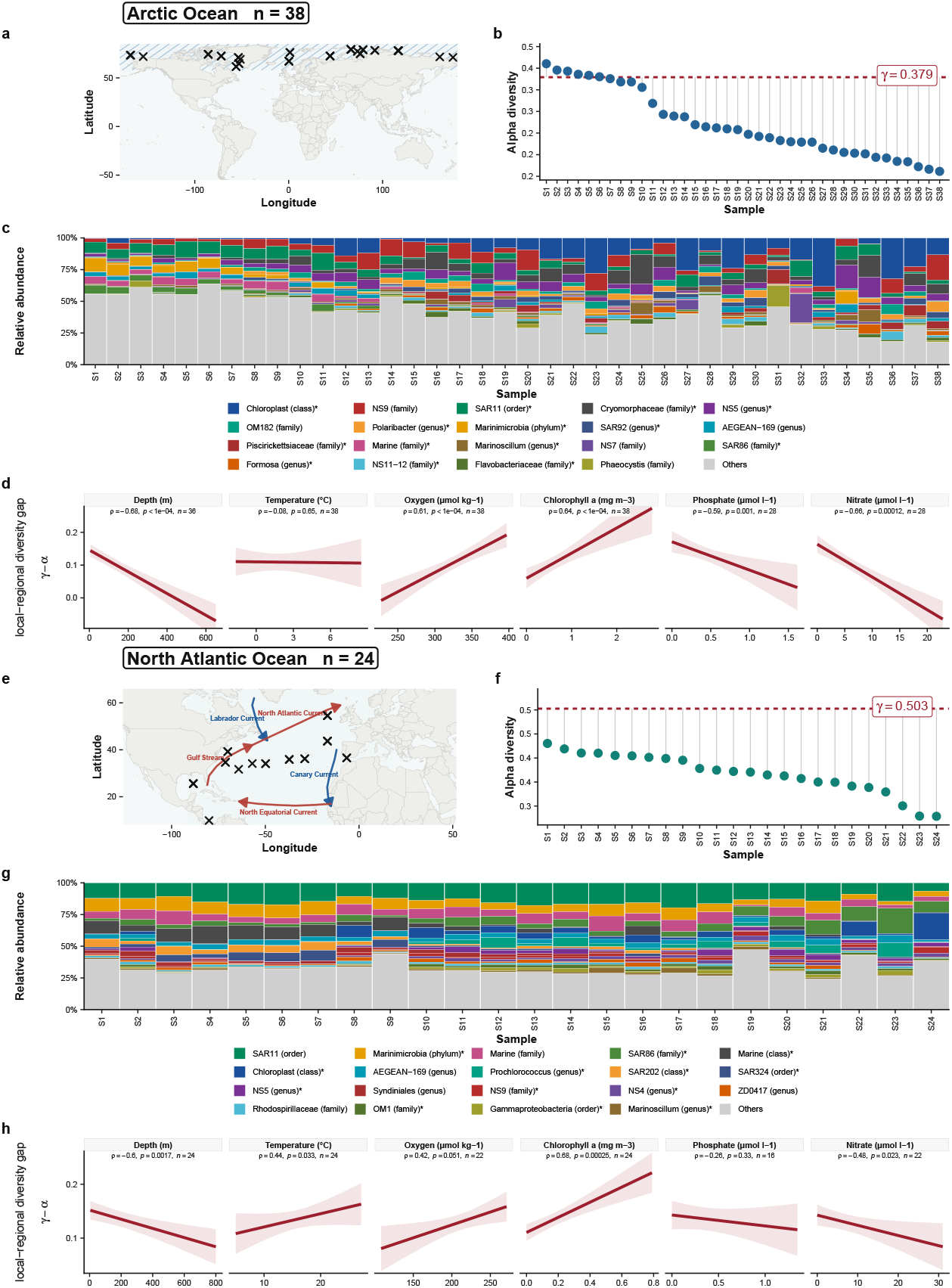
Concordant environmental associations of local–regional diversity coexist with region-specific taxon-level patterns in Arctic and North Atlantic waters. **a–d**, Arctic Ocean communities (*n* = 38). **e–h**, North Atlantic Ocean communities (*n* = 24). **a**,**e**, Sampling locations, shown as black crosses. Schematic warm and cold currents in **e** situate the North Atlantic stations within the major circulation pathways connecting subtropical, subpolar and Arctic waters. **b**,**f**, Sample-level HRIC alpha diversity calculated using `HRIC::SHalpha`. Samples are ordered by increasing signed local–regional diversity gap, defined as *γ − α*_*i*_. Dashed red lines show the diversity of the equal-community-weighted pooled regional composition, calculated using `HRIC::SHgamma`. **c**,**g**, Relative abundance profiles of the 19 taxa with the largest mean relative abundance in each region, with all remaining taxa grouped as Others. Bars follow the sample order in **b** and **f**. Asterisks identify taxa whose HRIC-transformed coordinates were associated with nitrate in univariable linear models at *P*_BH_ *<* 0.05, after Benjamini–Hochberg adjustment across the 19 tested taxa within each region. Nitrate was selected for the taxon-resolved panels because it had the largest number of adjusted associations among the six displayed environmental covariates. **d**,**h**, Signed local–regional diversity gap in relation to depth, temperature, oxygen, chlorophyll *a*, phosphate and nitrate. Red lines and shaded areas show ordinary least-squares visual summaries and 95% confidence intervals. Annotations report Spearman’s *ρ*, unadjusted two-sided *P* values and complete-case sample sizes.

Within each region, alpha diversity described diversity within individual local communities, whereas gamma diversity described diversity of the equal-community-weighted pooled regional composition. The Arctic ensemble had mean alpha diversity of 0.270 and gamma diversity of 0.379, yielding *β*_SH_ = 0.109; corresponding values in the North Atlantic were 0.368, 0.503 and 0.135. Because the two ensembles used different feature dictionaries, these absolute HRIC diversity values were interpreted within regions and not compared directly between regions. For sample-resolved analyses, we defined the signed local–regional diversity gap as *γ* − *α*_*i*_. Samples (Fig. 2 b,f) were ordered by this quantity so that scalar diversity organization could be compared with the corresponding taxonomic profiles.

Depth, chlorophyll *a* and nitrate showed concordant associations with the local– regional diversity gap in both regions. The gap declined with depth in the Arctic (Spearman’s *ρ* = −0.68) and North Atlantic (*ρ* = −0.60), increased with chlorophyll *a* (*ρ* = 0.64 and 0.68, respectively), and declined with nitrate (*ρ* = −0.66 and −0.48). Thus, expressed in terms of local diversity, alpha diversity tended to be higher relative to the corresponding gamma diversity in deeper and nitrate-richer waters and lower in chlorophyll-rich waters. Associations with temperature, oxygen and phosphate were less consistent between regions (Fig. 2 d,h). These patterns accord with the strong vertical organization previously reported for ocean microbial communities and with reported associations of nutrient supply and chlorophyll structure with plankton-microbiome diversity and composition [1, 20].

We next asked whether these concordant scalar associations were accompanied by shared taxon-level associations. Among the six environmental variables, nitrate had the largest number of Benjamini–Hochberg-adjusted associations with HRIC taxon coordinates and was therefore selected for an exploratory taxon-resolved follow-up. Among the 19 most abundant taxa in each region, 14 Arctic and 13 North Atlantic taxa were associated with nitrate at *P*_BH_ *<* 0.05. The overlap was only partial. Marinimicrobia showed positive coordinate associations with nitrate in both regions, whereas

*NS5*, chloroplast-assigned sequences and *Marinoscillum* showed negative associations in both; *SAR86* changed direction between the Arctic and North Atlantic. Other associated taxa differed between regions, with the Arctic pattern including several lineages associated with chlorophyll-rich upper-water communities and the North Atlantic pattern including lineages characteristic of deeper or energy-stratified waters [21–24]. Overall, nitrate was associated with coordinate variation across much of the displayed dominant assemblage, but the associated taxonomic pattern was region dependent.

Across the two regional ensembles, the local–regional diversity gap had concordant associations with major vertical environmental gradients, whereas the corresponding taxon-level associations were only partly shared. The analysis therefore separates two ecological statements that need not coincide: the same environmental gradients can be associated with similar scalar local–regional diversity patterns across regions while the associated taxa differ. By representing local alpha diversity, gamma diversity of the pooled regional composition, additive regional beta diversity and taxon-level compositional variation within the same framework, HRIC makes these scalar and taxonomic levels directly comparable without treating them as the same ecological quantity.

### 2.3 Auto-FMT is associated with earlier gut microbiota recovery after allo-HSCT

Allogeneic haematopoietic stem-cell transplantation (allo-HSCT) subjects the gut microbiota to conditioning, immunosuppression and intensive antibiotic exposure, often producing loss of alpha diversity and expansion of individual taxa that can dominate the community [25–27]. Autologous faecal microbiota transplantation (auto-FMT) aims to promote reconstitution by returning a patient’s pathogen-screened pre-treatment microbiota. Previous studies reported recovery of within-sample diversity and greater similarity to pre-treatment microbiota after autologous transfer [4, 28, 29]. We reanalysed the randomized allo-HSCT trial of Taur et al. [4] to ask whether these recovery dimensions could be separated within the same local–regional framework and related to their taxonomic contributions.

The evaluable randomized microbiome cohort comprised 25 patients, including 14 auto-FMT recipients and 11 controls, with 498 longitudinal stools. The primary longitudinal analysis contained 98 post-index stools from 24 patients over event days 1–60, with days 1–30 defining the primary early-recovery window. Each stool was treated as a local community. HRIC alpha diversity quantified within-stool diversity, whereas pairwise HRIC dissimilarity from each patient’s earliest observed pre-HSCT stool quantified return towards that personal reference composition. Local–regional summaries provided a complementary scale-resolved view: stools within clinical phases formed temporal ensembles for the patient-level trajectories in Fig. 3, whereas the landmark analysis (Fig. 4c) treated each randomized arm within the pre-index, early post-index and later post-index windows as a separate regional ensemble. Gamma diversity quantified the diversity of the corresponding pooled regional composition, and additive regional beta diversity related mean local alpha diversity to gamma diversity.

**Fig. 3.**
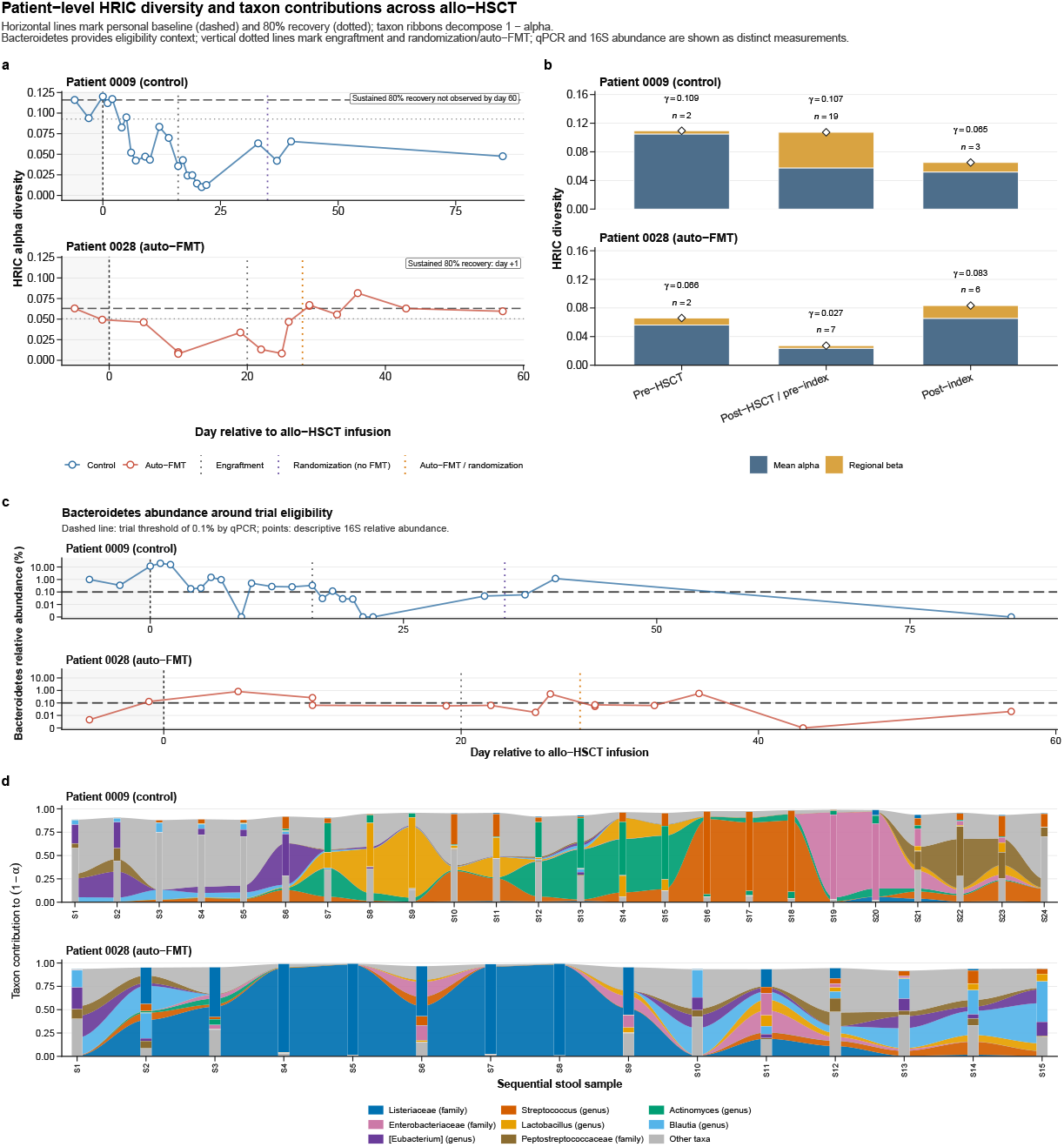
Patient-level HRIC trajectories resolve complementary dimensions of microbiota recovery after allo-HSCT. **a**, Longitudinal HRIC alpha diversity for patient 0009, assigned to control, and patient 0028, assigned to auto-FMT. Time is aligned to allo-HSCT infusion (clinical day 0). Vertical dotted lines mark neutrophil engraftment and randomization or auto-FMT. The horizontal dashed line denotes the earliest observed pre-HSCT alpha diversity for each patient, and the dotted line denotes 80% of that value. A sustained threshold crossing was confirmed at the next available sample. **b**, Within-patient HRIC decomposition before HSCT, after HSCT but before the index day and after the index day. Bars show mean alpha diversity and additive regional beta diversity; diamonds show gamma diversity of the equal-community-weighted pooled phase composition, with 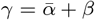. Values of *n* are stool samples. **c**, Bacteroidetes relative abundance measured by 16S rRNA gene sequencing. The horizontal line at 0.1% denotes the quantitative-PCR threshold used to determine trial eligibility and is shown only for clinical context; the threshold is not numerically interchangeable with the plotted 16S relative abundance. **d**, Additive taxon contributions to the complement of HRIC alpha diversity, 1 *− α*, across sequential stools. The eight taxa with the largest aggregate contributions across the two patients are displayed, with the remaining taxa combined as other taxa. Ribbon height is the taxon’s additive contribution to community unevenness. The two patients illustrate contrasting longitudinal recovery paths; randomized arm-level comparisons are shown in Fig. 4.

**Fig. 4.**
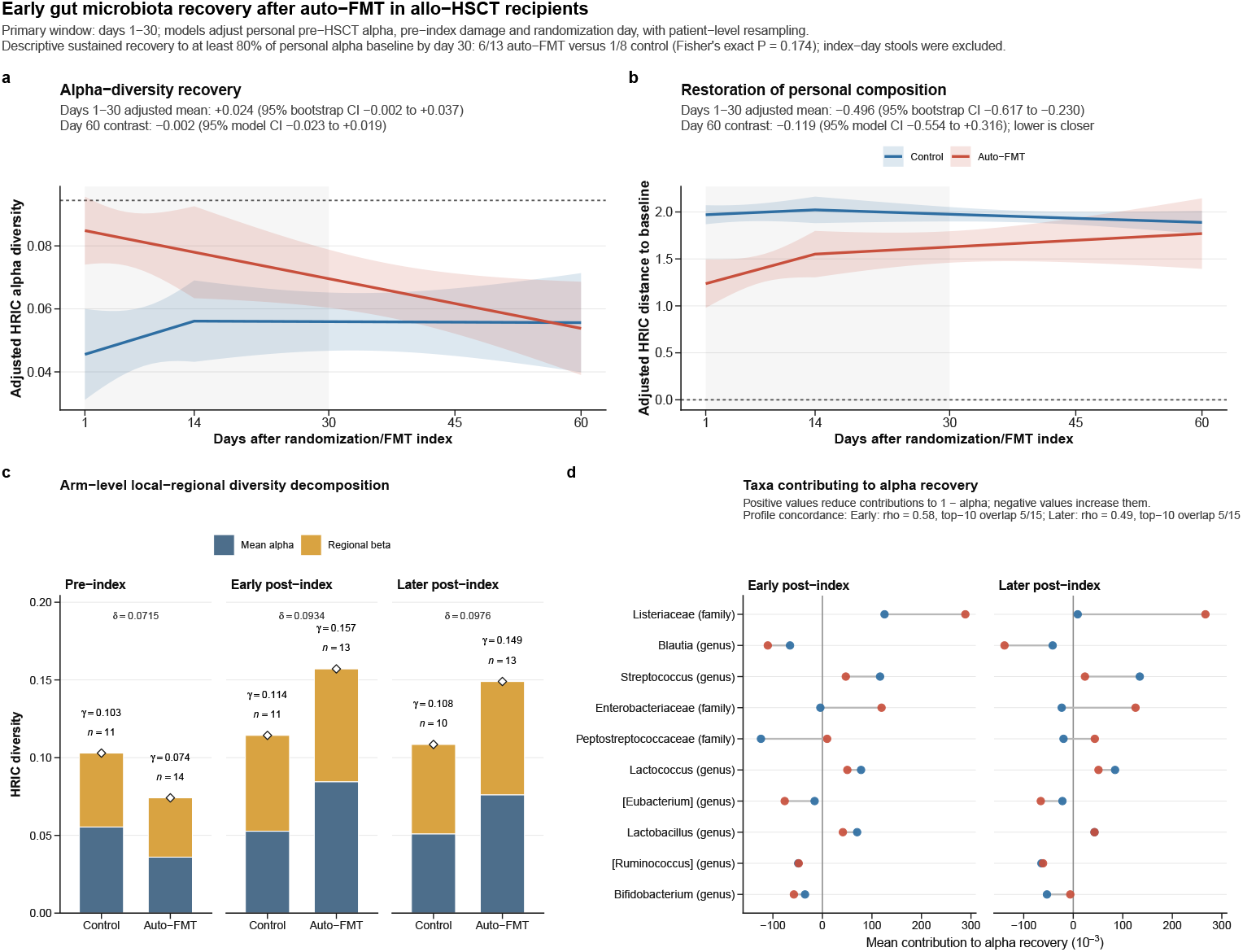
Auto-FMT is associated most clearly with earlier restoration of personal community composition. **a**, Adjusted HRIC alpha-diversity trajectories during event days 1–60. Fixed-effect predictions are from a mixed-effects model containing treatment arm, piecewise-linear event time (days 1–14 and 15–60), arm-by-time interactions, personal pre-HSCT alpha diversity, nearest pre-index alpha diversity, randomization day and a patient random intercept. The horizontal dashed line denotes mean personal pre-HSCT alpha diversity and the shaded vertical region denotes the primary days 1–30 window. The time-averaged auto-FMT-minus-control contrast was +0.0235 (95% stratified patient-bootstrap interval, *−* 0.0015 to +0.0373; *P* = 0.0668; 4,999 resamples). **b**, Adjusted pairwise HRIC dissimilarity from each patient’s earliest pre-HSCT sample; lower values indicate closer restoration of personal composition. Predictions are from a linear model containing treatment arm, piecewise-linear event time, arm-by-time interactions, personal pre-HSCT alpha diversity, pre-index HRIC dissimilarity and randomization day, with patient-clustered HC2 covariance. The horizontal dashed line at zero denotes exact return to the personal reference. The days 1–30 time-averaged auto-FMT-minus-control contrast was *−* 0.496 (95% stratified patient-bootstrap interval, *−*0.617 to *−*0.230; *P* = 4.0 10^*−*4^; 4,999 resamples). Curves and ribbons in **a** and **b** are fixed-effect predictions and pointwise model-based 95% confidence intervals; the time-averaged contrasts and their intervals were obtained from the stratified patient bootstrap. **c**, Arm-specific HRIC local–regional diversity decompositions at the nearest stool in the pre-index (days *−*28 to *−*1), early post-index (days 1–14) and later post-index (days 15–60) windows. Bars show mean alpha diversity and additive regional beta diversity; diamonds show gamma diversity of the arm-specific pooled landmark composition, and the mean of *γ − α*_*i*_ equals regional beta diversity. Values of *n* are patients; the *δ* annotations, calculated using `HRIC::SHdelta`, are separate from the local–regional diversity decomposition and report unnormalized treatment-arm-associated HRIC dispersion at each landmark. **d**, Taxon-level contributions to alpha-diversity recovery from the pre-index landmark to the early and later post-index landmarks. Positive values indicate a reduction in the taxon’s contribution to 1 *− α*. The evaluable randomized microbiome cohort comprised 25 patients (14 auto-FMT and 11 control); the primary longitudinal analysis contained 98 post-index stools from 24 patients.

The clearest randomized-arm difference concerned pairwise HRIC dissimilarity from the personal pre-HSCT composition (Fig. 4b). Averaged over event days 1–30, this adjusted dissimilarity was 0.496 units lower with auto-FMT than with control (95% stratified patient-bootstrap interval, −0.617 to −0.230; *P* = 4.0 × 10^*−*4^; 4,999 resamples). The contrast narrowed to −0.119 by day 60 (95% model-based CI, −0.554 to +0.316). Local alpha diversity showed a directionally similar but less precise early difference (Fig. 4 a): the days 1–30 adjusted auto-FMT-minus-control contrast was +0.0235 (95% bootstrap interval, −0.0015 to +0.0373; *P* = 0.0668) and was close to zero by day 60 (−0.0018). A corroborative event-time difference-in-differences analysis gave the same qualitative early separation for both outcomes (Supplementary Sections S5.6 and S5.9). Thus, the strongest early auto-FMT-associated pattern was closer return towards each patient’s pre-HSCT community composition, accompanied by a smaller and less precisely estimated difference in local diversity.

The landmark decompositions showed that these recovery dimensions did not move in parallel (Fig. 4 c). In the auto-FMT arm, higher post-index mean alpha diversity was accompanied by higher gamma diversity of the pooled landmark composition, and additive regional beta diversity did not simply contract as local diversity increased. Treatment-associated HRIC dispersion was also similar at the early and later post-index landmarks (*δ* = 0.0934 and 0.0976, respectively), compared with 0.0715 pre-index. These descriptive summaries therefore distinguish recovery of within-community diversity from changes in local–regional differentiation and from treatment-associated HRIC dispersion, rather than treating them as interchangeable measures of recovery.

Patient-level trajectories illustrated the heterogeneity underlying these arm-level summaries (Fig. 3). Patient 0028, assigned to auto-FMT, showed a sharp post-index rise in alpha diversity after a period of severe depletion, together with an increase in phase-level gamma diversity; patient 0009, assigned to control, passed through a more prolonged series of low-diversity states. The *Bacteroidetes* profiles provide clinical context for trial eligibility, while the HRIC taxon decomposition shows that similar scalar diversity values can arise from different community configurations. In patient 0028, the large pre-index contribution of *Listeriaceae* to 1 − *α* declined after auto-FMT, whereas patient 0009 moved through several taxon-dominated configurations during follow-up.

At the randomized-arm level, taxon contributions similarly showed that recovery of alpha diversity was compositionally heterogeneous (Fig. 4d). The largest early reductions in contributions to community unevenness in the auto-FMT arm involved *Listeriaceae* and *Enterobacteriaceae*, followed by smaller reductions for *Lactococcus, Streptococcus* and *Lactobacillus*. The contributing taxa overlapped only partly between treatment arms, consistent with individualized compositional trajectories rather than a single taxonomic recovery pattern. HRIC alpha was strongly concordant with the log inverse-Simpson index used in the original trial and with Pielou evenness, while localized disagreements between indices could be resolved through the taxon-level decomposition (Supplementary Sections S5.8 and S5.9).

Together, the randomized longitudinal and local–regional analyses identify a temporally concentrated auto-FMT-associated recovery pattern. During the first month, recipients were substantially closer to their personal pre-HSCT reference compositions, whereas the corresponding difference in local alpha diversity was smaller and less precise; both longitudinal contrasts attenuated by day 60. The landmark decompositions further show that recovery of local diversity does not imply a parallel reduction in local–regional differentiation. Auto-FMT is therefore associated most clearly with earlier personal compositional reconstitution, while local alpha diversity, local–regional differentiation and taxonomic composition capture complementary aspects of the recovery process.

## 3 Discussion

Microbiome diversity analysis inherits two ecological traditions often compressed within the alpha–beta vocabulary. Whittaker’s hierarchy defines alpha within local communities, gamma for the pooled regional composition and regional beta as local–regional differentiation [7]; Hill and Jost gave multiplicative regional beta an effective-number interpretation [8, 9]. In parallel, dissimilarity-based analysis developed from gradient ordination to variance partitioning, multivariate dispersion and association testing [10, 12, 30]. Our framework connects these traditions without treating regional beta diversity, pairwise dissimilarity and predictor-associated dispersion as synonyms: local and regional units are specified first, then quantified through a common representation.

HRIC provides this representation. Its square-root embedding and logarithmic map at the uniform composition produce finite, bounded Euclidean contrasts anchored in Fisher–Rao spherical geometry [16]. Simplex Hellinger alpha and gamma diversity quantify local and pooled compositions; gamma minus mean local alpha defines additive regional beta diversity, while the coordinates separately support pairwise dissimilarity and predictor-associated dispersion. HRIC alpha is a bounded monotonic reparameterization of the order-1*/*2 Hill number and therefore emphasizes low-abundance taxa more than Shannon or inverse Simpson diversity [8]. Hill numbers remain complementary when effective diversity across orders is the target; HRIC instead unifies sparse compositions, diversity partitions, taxon contributions and multivariate inference [8, 9].

Not every cohort constitutes a region. A regional ensemble must be prespecified on ecological or study-design grounds, with comparable local communities, a common feature dictionary and an explicit weighting rule. Direct dispersal or a shared source pool is not required, but cohort membership alone is insufficient when habitats, hosts, time windows or technical regimes are not jointly interpretable.

The ocean analysis illustrates why local–regional diversity and taxonomic compositional change are related but distinct. In both regional ensembles, the signed diversity gap had concordant associations with depth, chlorophyll *a* and nitrate, consistent with the vertical organization of global ocean microbiota [1, 2, 19]. Nitrate-associated taxa overlapped only partly: *Marinimicrobia, NS5, Marinoscillum* and chloroplast-assigned sequences showed shared directions, whereas *SAR86* changed direction and other lineages were region-specific. Thus, concordant diversity–environment associations need not imply shared taxon-level responses.

The auto-FMT analysis extends this distinction to host-associated communities through time. Post-index local and pooled regional diversity increased, but additive regional beta diversity did not simply contract as alpha recovered. Auto-FMT recipients were closer to their personal pre-HSCT compositions during days 1–30, whereas the alpha-diversity contrast was smaller and less precise; both attenuated by day 60. Contributors to alpha recovery also overlapped only partly between arms. The original trial established increased inverse-Simpson diversity and return towards pre-treatment composition after auto-FMT [4]; HRIC separates these outcomes as local diversity, local–regional differentiation and personal compositional recovery. The results support an earlier auto-FMT-associated return towards pre-HSCT composition, not a persistent diversity advantage, a distinction relevant to adverse outcomes associated with microbiota domination after allo-HSCT [25–27].

The framework targets compositional organization rather than all dimensions of microbial diversity. Incidence-based differentiation, phylogenetic replacement, functional change and alternative Hill orders remain complementary; occurrence-sensitive dissimilarities were more responsive to prevalence-only changes in simulations. HRIC also makes a deliberate geometric choice. Unlike Aitchison geometry, it does not preserve perturbation, powering or subcompositional coherence on the strictly positive simplex [13, 31]. The classical square-root/Hellinger embedding alone places compositions on a curved sphere, where ordinary coordinate-wise means, variances and regression summaries do not generally respect the manifold and Euclidean averages leave it. The logarithmic map instead supplies invertible coordinates in a flat tangent space while retaining radial Fisher–Rao geometry. Thus, radial distance from uniformity retains intrinsic information distance up to its conventional factor, while tangent coordinates support Euclidean statistical models [16]. The representation is intended for a fixed feature dictionary and accommodates observed zeros without replacement, but does not identify the observation process generating them. Sampling models that distinguish biological absence from depth-dependent non-detection are an important extension [32, 33], alongside alternative reference points, intrinsic pairwise models and weighted or phylogenetic feature spaces.

## 4 Methods

### Data sources and software

The ocean analysis used the publicly available Tara Oceans prokaryote-enriched metagenomic 16S/18S rRNA miTAG abundance tables described by Salazar et al. [2]. Profiles for 180 samples are available through the study companion site (https://www.ocean-microbiome.org/) and EBI BioStudies accession `S-BSST297`; matched sample and environmental metadata are available in Tables W1–W6 and through PANGAEA (doi:10.1594/PANGAEA.875582) [17]. The auto-FMT analysis used the processed 16S rRNA gene feature table and sample metadata from Qiita study `12277` (https://qiita.ucsd.edu/study/description/12277), corresponding to the randomized auto-FMT trial NCT02269150 reported by Taur et al. [4]; after removal of empty samples and globally absent features, the dataset contained 498 stools from 25 evaluable randomized patients (14 auto-FMT and 11 control) and 6,647 features. Analyses were performed in R 4.5.0 using `HRIC` 0.1.0, `phyloseq` 1.52.0, `lme4` 2.0.1, `sandwich` 3.1.1, `ggplot2` 4.0.3, `dplyr` 1.2.1, `tidyr` 1.3.2 and `patchwork` 1.3.2; base-R `stats` functions were used for linear models, correlations, *t*-tests, Fisher’s exact tests and multiplicity adjustment, and random seeds were fixed before resampling and permutation procedures. Data filtering, metadata processing and analysis-specific implementation are described in Supplementary Note 5.

### HRIC representation and taxon contributions

All HRIC transformations and diversity summaries followed the definitions in the Results and Supplementary Notes 2 and 3. The feature dictionary was held fixed for every comparison assigned a common HRIC scale: all 6,647 gut features were retained across gut samples, whereas each ocean region used its own region-wide set of observed features. A taxon’s additive contribution to the complement of alpha diversity was its normalized squared HRIC coordinate, and contributions summed to 1 − *α* within each sample. Contributions were aggregated after assigning features to the deepest available taxonomic label; they quantify the taxonomic structure of HRIC diversity rather than relative abundance. Computational details and the correspondence between mathematical quantities and package functions are provided in Supplementary Note 4.

### Taxon-level association analyses

Taxon-level associations were analysed in HRIC coordinate space. For a two-level comparison, each taxon’s coordinate was compared between groups using Welch’s two-sample *t*-test (`stats::t.test`, unequal variances). For a continuous covariate, a separate univariable linear model was fitted to each taxon coordinate using `stats::lm`, and the two-sided coefficient test assessed a zero slope. Benjamini–Hochberg adjustment was applied separately within each prespecified set of taxa and covariate. Condition-associated HRIC dispersion and taxon-level association testing address different scales: `SHdelta` summarizes the weighted separation of condition-specific centroids in the full HRIC coordinate space, whereas taxon-level models localize coordinate associations to individual taxa.

### Environmental association analyses

Associations between the signed local–regional diversity gap and continuous covariates were assessed on complete cases using two-sided Spearman rank correlation (`stats::cor.test`, `method=“spearman”`, `exact=FALSE`). Ordinary least-squares lines and 95% confidence bands in the correlation panels were visual summaries and were not used to calculate the reported Spearman statistics. Unless stated otherwise, tests were two-sided. Taxon-level test families were multiplicity-adjusted as described above; the ocean diversity-gap–environment correlations were exploratory and their reported *P* values were unadjusted. The simulation design and PERMANOVA correspondence are detailed in Supplementary Note 3, and the ocean and auto-FMT implementations, longitudinal models, resampling procedures, sensitivity analyses and conventional-index comparisons are detailed in Supplementary Note 5.

### Auto-FMT longitudinal analyses

For randomized comparisons, event time was defined as stool-collection day minus randomization day, such that event day 0 denoted auto-FMT in the treatment arm and randomization in controls; exact event-day-0 stools were excluded because their collection order relative to auto-FMT was unavailable. Each patient’s earliest observed pre-HSCT stool served as the personal compositional reference. The primary longitudinal dataset comprised 98 stools from 24 patients during event days 1–60, and the primary estimand was the adjusted auto-FMT-minus-control contrast averaged over days 1–30. HRIC alpha diversity was analysed with a piecewise-linear mixed-effects model containing treatment arm, event-time terms for days 1–14 and 15–60, arm-by-time interactions, personal pre-HSCT alpha diversity, nearest pre-index alpha diversity, randomization day and a patient random intercept. Because the corresponding random-intercept variance for pairwise HRIC dissimilarity from the personal reference was estimated as zero, this dissimilarity outcome was analysed by ordinary least squares with the same event-time structure, personal pre-HSCT alpha diversity, pre-index HRIC dissimilarity and randomization day, using patient-clustered HC2 covariance. The days 1–30 estimand was obtained by trapezoidal integration of the daily fixed-effect contrast curve, with uncertainty estimated from 4,999 non-parametric patient-level bootstrap resamples stratified by randomized arm. Landmark local–regional diversity decompositions, treatment-arm-associated HRIC dispersion and taxon-contribution summaries were descriptive. Full model specifications, landmark definitions, sensitivity analyses and corroborative recovery analyses are provided in Supplementary Note 5.

## Declarations

The authors declare no competing interests.

## Data availability

The Tara Oceans prokaryote-enriched miTAG profiles analysed in this study are available through the study companion site (https://www.ocean-microbiome.org/) and EBI BioStudies under accession `S-BSST297`; matched environmental metadata are available through PANGAEA (https://doi.org/10.1594/PANGAEA.875582). The processed auto-FMT feature table and sample metadata are available from Qiita study `12277` (https://qiita.ucsd.edu/study/description/12277).

## Code availability

All code required to reproduce the analyses, together with the processed input data, is publicly available at https://github.com/yiqianomics/HRIC. The repository contains the HRIC R package and the code used for all diversity, dissimilarity, dispersion and longitudinal analyses reported here. Documentation, tutorials and additional information about HRIC are available at the project website: https://yiqianzhang.com/hric/.

## Ethics Statement

This study is a secondary analysis of previously published, publicly available datasets and did not involve the collection of new data from human participants or animals. The ocean analysis used publicly available environmental metagenomic profiles and physicochemical metadata from the Tara Oceans expedition, which involve no human or animal subjects. The gut microbiota analysis used the de-identified, publicly available 16S rRNA gene feature table and sample metadata deposited in Qiita (study 12277) from the randomized trial NCT02269150. That trial was approved by the institutional review board of the originating institution, and written informed consent was obtained from all participants by the original investigators; the present authors had no access to identifiable participant information and no contact with participants. The authors’ institutional review board determined that this secondary analysis of de-identified public data does not constitute human-subjects research and confirmed that it is exempt from review.

## Author contributions

YZ: Data curation, Software, Formal analysis, Visualization, Validation, Investigation, Methodology, Writing–original draft, Writing–review&editing; ZZ: Conceptualization, Resources, Supervision, Project administration, Validation, Investigation, Methodology, Writing–original draft, Writing–review&editing.

## References

[1] Sunagawa, S., et al.: Structure and function of the global ocean microbiome. Science 348(6237), 1261359 (2015) 10.1126/science.1261359

[2] Salazar, G., et al.: Gene expression changes and community turnover differentially shape the global ocean metatranscriptome. Cell 179(5), 1068–108321 (2019) 10.1016/j.cell.2019.10.014

[3] The Human Microbiome Project Consortium: Structure, function and diversity of the healthy human microbiome. Nature 486(7402), 207–214 (2012) 10.1038/nature11234

[4] Taur, Y., Coyte, K., Schluter, J., Robilotti, E., Figueroa, C., Gjonbalaj, M., Littmann, E.R., Ling, L., Miller, L., Gyaltshen, Y., Fontana, E., Morjaria, S., Gyurkocza, B., Perales, M.-A., Castro-Malaspina, H., Tamari, R., Ponce, D., Koehne, G., Barker, J., Jakubowski, A., Papadopoulos, E., Dahi, P., Sauter, C., Shaffer, B., Young, J.W., Peled, J.U., Meagher, R.C., Jenq, R.R., Brink, M.R.M., Giralt, S.A., Pamer, E.G., Xavier, J.B.: Reconstitution of the gut microbiota of antibiotic-treated patients by autologous fecal microbiota transplant. Science Translational Medicine 10(460), 9489 (2018) 10.1126/scitranslmed.aap9489

[5] Caporaso, J.G., Kuczynski, J., Stombaugh, J., Bittinger, K., Bushman, F.D., Costello, E.K., Fierer, N., Gonzalez Peña, A., Goodrich, J.K., Gordon, J.I., Huttley, G.A., Kelley, S.T., Knights, D., Koenig, J.E., Ley, R.E., Lozupone, C.A., McDonald, D., Muegge, B.D., Pirrung, M., Reeder, J., Sevinsky, J.R., Turnbaugh, P.J., Walters, W.A., Widmann, J., Yatsunenko, T., Zaneveld, J., Knight, R.: QIIME allows analysis of high-throughput community sequencing data. Nature Methods 7(5), 335–336 (2010) 10.1038/nmeth.f.303

[6] Lozupone, C., Knight, R.: UniFrac: A new phylogenetic method for comparing microbial communities. Applied and Environmental Microbiology 71(12), 8228–8235 (2005) 10.1128/AEM.71.12.8228-8235.2005

[7] Whittaker, R.H.: Evolution and measurement of species diversity. Taxon 21(2–3), 213–251 (1972) 10.2307/1218190

[8] Hill, M.O.: Diversity and evenness: A unifying notation and its consequences. Ecology 54(2), 427–432 (1973) 10.2307/1934352

[9] Jost, L.: Partitioning diversity into independent alpha and beta components. Ecology 88(10), 2427–2439 (2007) 10.1890/06-1736.1

[10] Bray, J.R., Curtis, J.T.: An ordination of the upland forest communities of southern wisconsin. Ecological Monographs 27(4), 325–349 (1957) 10.2307/1942268

[11] Anderson, M.J., Crist, T.O., Chase, J.M., Vellend, M., Inouye, B.D., Freestone, A.L., Sanders, N.J., Cornell, H.V., Comita, L.S., Davies, K.F., Harrison, S.P., Kraft, N.J.B., Stegen, J.C., Swenson, N.G.: Navigating the multiple meanings of beta diversity: A roadmap for the practicing ecologist. Ecology Letters 14(1), 19–28 (2011) 10.1111/j.1461-0248.2010.01552.x

[12] Legendre, P., De Cáceres, M.: Beta diversity as the variance of community data: Dissimilarity coefficients and partitioning. Ecology Letters 16(8), 951–963 (2013) 10.1111/ele.12141

[13] Aitchison, J.: The statistical analysis of compositional data. Journal of the Royal Statistical Society: Series B (Methodological) 44(2), 139–177 (1982) 10.1111/j.2517-6161.1982.tb01195.x

[14] Lubbe, S., Filzmoser, P., Templ, M.: Comparison of zero replacement strategies for compositional data with large numbers of zeros. Chemometrics and Intelligent Laboratory Systems 210, 104248 (2021) 10.1016/j.chemolab.2021.104248

[15] Lande, R.: Statistics and partitioning of species diversity, and similarity among multiple communities. Oikos 76(1), 5–13 (1996) 10.2307/3545743

[16] Rao, C.R.: Information and the accuracy attainable in the estimation of statistical parameters. Bulletin of the Calcutta Mathematical Society 37(3), 81–91 (1945)

[17] Pesant, S., et al.: Open science resources for the discovery and analysis of Tara Oceans data. Scientific Data 2, 150023 (2015) 10.1038/sdata.2015.23

[18] Wietz, M., Bienhold, C., Metfies, K., Torres-Valdés, S., Appen, W.-J., Salter, I., Boetius, A.: The polar night shift: Seasonal dynamics and drivers of arctic ocean microbiomes revealed by autonomous sampling. ISME Communications 1(1), 76 (2021) 10.1038/s43705-021-00074-4

[19] Richter, D.J., et al.: Genomic evidence for global ocean plankton biogeography shaped by large-scale current systems. eLife 11, 78129 (2022) 10.7554/eLife.78129

[20] James, C.C., Barton, A.D., Allen, L.Z., Lampe, R.H., Rabines, A., Schulberg, A., Zheng, H., Goericke, R., Goodwin, K.D., Allen, A.E.: Influence of nutrient supply on plankton microbiome biodiversity and distribution in a coastal upwelling region. Nature Communications 13(1), 2448 (2022) 10.1038/s41467-022-30139-4

[21] Teeling, H., Fuchs, B.M., Bennke, C.M., Krüger, K., Chafee, M., Kappelmann, L., Reintjes, G., Waldmann, J., Quast, C., Glöckner, F.O., Lucas, J., Wichels, A., Gerdts, G., Wiltshire, K.H., Amann, R.I.: Recurring patterns in bacterioplankton dynamics during coastal spring algae blooms. eLife 5, 11888 (2016) 10.7554/eLife.11888

[22] Mehrshad, M., Rodriguez-Valera, F., Amoozegar, M.A., López-García, P., Ghai, R.: The enigmatic SAR202 cluster up close: Shedding light on a globally distributed dark ocean lineage involved in sulfur cycling. The ISME Journal 12(3), 655–668 (2018) 10.1038/s41396-017-0009-5

[23] Sheik, C.S., Jain, S., Dick, G.J.: Metabolic flexibility of enigmatic SAR324 revealed through metagenomics and metatranscriptomics. Environmental Microbiology 16(1), 304–317 (2014) 10.1111/1462-2920.12165

[24] Hawley, A.K., Nobu, M.K., Wright, J.J., Durno, W.E., Morgan-Lang, C., Sage, B., Schwientek, P., Swan, B.K., Rinke, C., Torres-Beltrán, M., Mewis, K., Liu, W.-T., Stepanauskas, R., Woyke, T., Hallam, S.J.: Diverse marinimicrobia bacteria may mediate coupled biogeochemical cycles along eco-thermodynamic gradients. Nature Communications 8(1), 1507 (2017) 10.1038/s41467-017-01376-9

[25] Taur, Y., Xavier, J.B., Lipuma, L., Ubeda, C., Goldberg, J., Gobourne, A., Lee, Y.J., Dubin, K.A., Socci, N.D., Viale, A., Perales, M.-A., Jenq, R.R., Brink, M.R.M., Pamer, E.G.: Intestinal domination and the risk of bacteremia in patients undergoing allogeneic hematopoietic stem cell transplantation. Clinical Infectious Diseases 55(7), 905–914 (2012) 10.1093/cid/cis580

[26] Shono, Y., et al.: Increased GVHD-related mortality with broad-spectrum antibiotic use after allogeneic hematopoietic stem cell transplantation in human patients and mice. Science Translational Medicine 8(339), 339–71 (2016) 10.1126/scitranslmed.aaf2311

[27] Peled, J.U., et al.: Microbiota as predictor of mortality in allogeneic hematopoietic-cell transplantation. The New England Journal of Medicine 382(9), 822–834 (2020) 10.1056/NEJMoa1900623

[28] Suez, J., Zmora, N., Zilberman-Schapira, G., Mor, U., Dori-Bachash, M., Bashiardes, S., Zur, M., Regev-Lehavi, D., Ben-Zeev Brik, R., Federici, S., Horn, M., Cohen, Y., Moor, A.E., Zeevi, D., Korem, T., Kotler, E., Harmelin, A., Itzkovitz, S., Maharshak, N., Shibolet, O., Pevsner-Fischer, M., Shapiro, H., Sharon, I., Halpern, Z., Segal, E., Elinav, E.: Post-antibiotic gut mucosal microbiome reconstitution is impaired by probiotics and improved by autologous FMT. Cell 174(6), 1406–142316 (2018) 10.1016/j.cell.2018.08.047

[29] Malard, F., Vekhoff, A., Lapusan, S., Isnard, F., D’incan-Corda, E., Rey, J., Saillard, C., Thomas, X., Ducastelle-Lepretre, S., Paubelle, E., Larcher, M.-V., Rocher, C., Recher, C., Tavitian, S., Bertoli, S., Michallet, A.-S., Gilis, L., Peterlin, P., Chevallier, P., Nguyen, S., Plantamura, E., Boucinha, L., Gasc, C., Michallet, M., Dore, J., Legrand, O., Mohty, M.: Gut microbiota diversity after autologous fecal microbiota transfer in acute myeloid leukemia patients. Nature Communications 12(1), 3084 (2021) 10.1038/s41467-021-23376-6

[30] Anderson, M.J., Ellingsen, K.E., McArdle, B.H.: Multivariate dispersion as a measure of beta diversity. Ecology Letters 9(6), 683–693 (2006) 10.1111/j.1461-0248.2006.00926.x

[31] Egozcue, J.J., Pawlowsky-Glahn, V., Mateu-Figueras, G., Barceló-Vidal, C.: Isometric logratio transformations for compositional data analysis. Mathematical Geology 35(3), 279–300 (2003) 10.1023/A:1023818214614

[32] Kaul, A., Davidov, O., Peddada, S.D.: Structural zeros in high-dimensional data with applications to microbiome studies. Biostatistics 18(3), 422–433 (2017) 10.1093/biostatistics/kxw053

[33] Chan, L.S., Li, G.: Zero is not absence: censoring-based differential abundance analysis for microbiome data. Bioinformatics 40(2), 071 (2024) 10.1093/bioinformatics/btae071

