## Supplementary Information for "Rebuilding microbiome diversity theory on the closed simplex"

### Supplementary Information for
### "Rebuilding microbiome diversity theory on the closed simplex"

Yiqian Zhang[1,2] Zihan Zhu[3,*]

[1]Interdisciplinary Ph.D. Program in Biostatistics, The Ohio State University, 1841 Neil Ave., Columbus, OH 43210, USA
[2]Department of Statistics, The Ohio State University, 1958 Neil Ave., Columbus, OH 43210, USA
[3]Department of Biostatistics, Yale School of Public Health, New Haven, Connecticut, USA

**Abstract**

This Supplementary Information develops the ecological and mathematical foundations of Hellinger–Riemann intrinsic coordinates (HRIC) and the associated Simplex Hellinger diversity framework. It reviews local–regional diversity partitioning and the development of dissimilarity-based community analysis; defines HRIC on the closed simplex; establishes zero compatibility, Euclidean contrast structure, boundedness, invertibility, boundary behaviour and the local relationship with log-ratio coordinates; and derives Simplex Hellinger alpha and gamma diversity, additive regional beta diversity, pairwise dissimilarity and predictor-associated HRIC dispersion from the same representation. It also provides the PERMANOVA correspondence, complete simulation design, worked example, computational algorithm, software mapping and statistical implementation for the ocean and auto-FMT analyses. All quantities are defined on a fixed feature dictionary, and all notation and equation references are self-contained.

### Supplementary Note 1: Historical foundations of local–regional diversity and dissimilarity analysis

#### S1.1 Additive and multiplicative local–regional partitions

Whittaker's local–regional framework separates diversity within local communities from the compositional differentiation through which those communities form a higher-level ensemble [1]. Two partitioning architectures subsequently became prominent. Additive partitioning defines regional beta diversity as the excess of gamma diversity over mean local alpha diversity,

$$\beta_{\text{add}} = \gamma - \bar{\alpha}, \tag{S1}$$

and is natural when the scientific target is an increment measured in the same units as alpha and gamma diversity [2]. Multiplicative partitioning defines

$$^{q}D_{\beta} = \frac{^{q}D_{\gamma}}{^{q}D_{\alpha}}, \tag{S2}$$

where the diversities are expressed as Hill numbers, or effective numbers of species [3, 4]. Jost showed that this effective-number formulation yields alpha and regional beta components without hidden dependence on one another and gives multiplicative regional beta diversity the interpretation of an effective number of compositionally distinct communities.

For three local communities containing three species each and five species collectively, the richness ratio $\gamma/\bar{\alpha} = 5/3$ states that regional richness is 5/3 times mean local richness; the effective-community interpretation follows from the later Hill–Jost framework rather than from the numerical ratio alone. If the three communities are uniform over their observed species, their order-2 Hill alpha diversity is $^{2}D_{\alpha} = 3$. The equally weighted pooled relative-abundance composition has $^{2}D_{\gamma} = 81/17$, giving $^{2}D_{\beta} = 27/17$. The pooled ensemble therefore has the same order-2 diversity as approximately 1.59 completely distinct communities of effective size 3. Additive and multiplicative regional beta diversity are interpreted within their respective architectures rather than as interchangeable transformations of a universal beta component.

#### S1.2 From local–regional differentiation to dissimilarity-based analysis

The decomposition and dissimilarity traditions emerged as related strands of the same ecological problem. Bray and Curtis developed polar ordination by arranging forest stands according to pairwise community resemblance, turning compositional differences into positions along ecological gradients [5]. Whittaker used the term beta diversity broadly, encompassing both the relation between local and regional diversity and community differentiation along gradients; he explicitly discussed dissimilarity between pairs of communities as a measure of beta differentiation [1]. Gauch and Whittaker subsequently compared ordination procedures on simulated coenoclines and showed that beta diversity and the choice of resemblance coefficient both influence ordination performance [6]. In this early literature, beta diversity, ecological distance and gradient length were closely connected but were not reduced to one universal index.

The proliferation of pairwise coefficients made the ecological target increasingly dependent on index choice. Koleff et al. reviewed 24 presence–absence beta-diversity measures and showed that they emphasize different combinations of shared species, richness gradients, species gains and species losses [7]. Their distinction between broad-sense measures, which include differences arising from richness gradients, and narrow-sense measures, which emphasize compositional replacement, demonstrated that numerical values labelled beta diversity can encode different ecological phenomena.

Tuomisto later organized these meanings by separating beta diversity defined as a function of alpha and gamma from measures of compositional heterogeneity, similarity, nestedness, gradient length and spatial variation [8].

A variance-based strand connected beta diversity directly to multivariate analysis. Legendre et al. defined the variance of a suitably represented community table as one measure of beta diversity and used canonical ordination to partition that variation among environmental and spatial components [9]. Anderson et al. represented beta diversity as multivariate dispersion, measured by the average dissimilarity of sampling units from a group centroid, and developed permutation procedures for comparing dispersion among groups [10]. Ricotta and Burrascano extended tests of beta-diversity differences to asymmetric plot-to-plot dissimilarities [11]. Legendre and De Cáceres subsequently showed that total compositional variance can be recovered from dissimilarity matrices satisfying appropriate properties and partitioned into species contributions, local contributions, among-group components, ordination axes, spatial scales and explanatory fractions [12].

Distance-based regression and PERMANOVA shifted the emphasis from describing total compositional heterogeneity to testing whether a specified predictor explains part of that heterogeneity [13, 14]. Microbiome kernel methods such as MiRKAT apply the same principle after converting a community dissimilarity into a similarity kernel [15]. These methods answer an association question: whether an environmental, host or experimental variable is aligned with multivariate community variation under the selected representation.

This progression supplied increasingly sophisticated indices, decompositions and tests, but the developments were usually constructed around a selected coefficient or transformed community table. Even variance-based approaches require the analyst to choose a representation or dissimilarity with suitable properties. They do not automatically require within-community diversity, diversity of the pooled regional composition, regional beta diversity, pairwise dissimilarity and predictor-associated HRIC dispersion to inherit one common representation of composition. The HRIC framework addresses this synthesis problem for sparse microbiome data by deriving all of these quantities from one Euclidean coordinate map of the closed simplex.

### Supplementary Note 2: Closed-simplex geometry and Hellinger–Riemann intrinsic coordinates

#### S2.1 Compositions, feature dictionaries and regional ensembles

Let $\boldsymbol{x} = (x_1, \ldots, x_p)^\mathsf{T} \in \mathbb{R}^p_{\geq 0}$ be a non-negative abundance vector with positive total. Its closure is

$$\boldsymbol{\pi} = \mathcal{C}(\boldsymbol{x}) = \frac{\boldsymbol{x}}{\sum_{j=1}^{p} x_j}. \tag{S3}$$

The resulting composition belongs to the closed simplex

$$\overline{\mathcal{S}}^{p-1} = \left\{ \boldsymbol{\pi} \in [0,1]^p : \sum_{j=1}^{p} \pi_j = 1 \right\}. \tag{S4}$$

Its relative interior is the open simplex

$$\mathcal{S}_+^{p-1} = \left\{ \boldsymbol{\pi} \in (0,1)^p : \sum_{j=1}^{p} \pi_j = 1 \right\}.$$

Exact zeros are ordinary boundary points of $\overline{\mathcal{S}}^{p-1}$. The uniform composition is

$$\boldsymbol{\pi}_0 = \left(p^{-1}, \dots, p^{-1}\right)^{\mathsf{T}}. \tag{S5}$$

Closure removes the arbitrary total because $\mathcal{C}(b\boldsymbol{x}) = \mathcal{C}(\boldsymbol{x})$ for every $b > 0$. An all-zero vector has no composition and must be removed or resolved as an assay failure before analysis.

All quantities are defined relative to a fixed dictionary of $p$ labelled features. Local zeros are retained, but a feature absent from every community in an analysis may be removed before the dictionary is fixed. Changing the dictionary changes $p$, the uniform reference $\boldsymbol{\pi}_0$, the normalizing constant introduced below and, in general, every HRIC coordinate. Comparisons assigned a common numerical scale must therefore use the same features in the same order.

A regional ensemble is an analytically prespecified collection of local compositions on this common dictionary for which pooling has a clear ecological interpretation. Its definition requires an explicit weighting rule and a study context in which the pooled composition represents the higher-level ensemble. It does not require a specified dispersal process or a demonstrably shared source pool. Material nuisance heterogeneity can be addressed through restriction, stratification or covariate adjustment, while the environmental, host or experimental factor of interest is analysed within the prespecified ensemble.

Let $w_i \geq 0$, with $\sum_{i=1}^{n} w_i = 1$, denote the weights assigned to the local communities, and write

$$\boldsymbol{y}_i = \sqrt{\boldsymbol{\pi}_i} \tag{S6}$$

for the element-wise square-root composition. The weighted Hellinger pool is

$$\bar{\boldsymbol{y}}_w = \sum_{i=1}^{n} w_i \boldsymbol{y}_i, \qquad \boldsymbol{y}_\gamma = \frac{\bar{\boldsymbol{y}}_w}{\|\bar{\boldsymbol{y}}_w\|_2}, \qquad \boldsymbol{\pi}_\gamma = \boldsymbol{y}_\gamma^{\circ 2}, \tag{S7}$$

where $^{\circ 2}$ denotes element-wise squaring. Normalization returns the weighted mean direction to the unit sphere, and element-wise squaring produces the pooled regional composition on the simplex. Equal weights, $w_i = 1/n$, give every local community the same influence irrespective of sequencing depth and are the package default used in the reported analyses.

#### S2.2 Square-root embedding and the contrast tangent space

Define the element-wise square-root map

$$\boldsymbol{y}(\boldsymbol{\pi}) = \sqrt{\boldsymbol{\pi}} = \left(\sqrt{\pi_1}, \dots, \sqrt{\pi_p}\right)^{\mathsf{T}}. \tag{S8}$$

Because $\sum_{j=1}^{p} \pi_j = 1$, $\|\boldsymbol{y}(\boldsymbol{\pi})\|_2 = 1$, so $\boldsymbol{y}$ maps $\overline{\mathcal{S}}^{p-1}$ bijectively to the non-negative orthant of the unit sphere $\mathbb{S}^{p-1} \subset \mathbb{R}^p$. The uniform composition maps to

$$\boldsymbol{y}_0 = \boldsymbol{y}(\boldsymbol{\pi}_0) = p^{-1/2}\mathbf{1}. \tag{S9}$$

For two compositions $\boldsymbol{\pi}$ and $\boldsymbol{\pi}'$,

$$\langle \boldsymbol{y}(\boldsymbol{\pi}), \boldsymbol{y}(\boldsymbol{\pi}') \rangle = \sum_{j=1}^{p} \sqrt{\pi_j \pi_j'} \tag{S10}$$

is the Bhattacharyya coefficient. Their square-root chordal distance satisfies

$$\|\boldsymbol{y}(\boldsymbol{\pi}) - \boldsymbol{y}(\boldsymbol{\pi}')\|_2^2 = 2\left[1 - \langle \boldsymbol{y}(\boldsymbol{\pi}), \boldsymbol{y}(\boldsymbol{\pi}') \rangle\right], \tag{S11}$$

and is $\sqrt{2}$ times the conventional Hellinger distance. The great-circle angle is

$$\theta(\boldsymbol{\pi}, \boldsymbol{\pi}') = \arccos \left\langle \boldsymbol{y}(\boldsymbol{\pi}), \boldsymbol{y}(\boldsymbol{\pi}') \right\rangle. \tag{S12}$$

Under the convention in which the multinomial Fisher–Rao line element is $\sum_{j=1}^{p}(d\pi_j)^2/\pi_j$, the Fisher–Rao distance is $2\theta(\boldsymbol{\pi}, \boldsymbol{\pi}')$; HRIC uses the angle itself [16]. The square-root map is continuous on the closed simplex. In particular, $\sqrt{\pi_j} \to 0$ as $\pi_j \to 0$, while the derivative $1/(2\sqrt{\pi_j})$ retains sensitivity near the boundary without producing a divergence in the mapped value.

The tangent space to the unit sphere at $\boldsymbol{y}_0$ is

$$T_{\boldsymbol{y}_0}\mathbb{S}^{p-1} = \left\{\boldsymbol{v} \in \mathbb{R}^p : \boldsymbol{v}^{\mathsf{T}}\boldsymbol{y}_0 = 0\right\} = \left\{\boldsymbol{v} \in \mathbb{R}^p : \sum_{j=1}^{p} v_j = 0\right\} =: \mathcal{H}_p. \tag{S13}$$

It is the $(p-1)$-dimensional zero-sum contrast subspace. For $\boldsymbol{\pi} \in \overline{\mathcal{S}}^{p-1}$, define

$$c(\boldsymbol{\pi}) = \langle \boldsymbol{y}(\boldsymbol{\pi}), \boldsymbol{y}_0 \rangle = \frac{1}{\sqrt{p}} \sum_{j=1}^{p} \sqrt{\pi_j}, \qquad s(\boldsymbol{\pi}) = \sqrt{1 - c(\boldsymbol{\pi})^2}, \qquad \theta(\boldsymbol{\pi}) = \arccos c(\boldsymbol{\pi}) = \arcsin s(\boldsymbol{\pi}). \tag{S14}$$

Because $\boldsymbol{y}(\boldsymbol{\pi})$ and $\boldsymbol{y}_0$ lie in the non-negative orthant, $\theta(\boldsymbol{\pi}) \in [0, \pi/2)$.

##### S2.3 Definition and logarithmic-map representation

**Definition S2.1** (Hellinger–Riemann intrinsic coordinates)**.** For $\boldsymbol{\pi} \in \overline{\mathcal{S}}^{p-1}$, define

$$\mathrm{HRIC}(\boldsymbol{\pi}) = \mathrm{Log}_{\boldsymbol{y}_0}\{\boldsymbol{y}(\boldsymbol{\pi})\} = \begin{cases} \dfrac{\theta(\boldsymbol{\pi})}{\sin\{\theta(\boldsymbol{\pi})\}}\left[\boldsymbol{y}(\boldsymbol{\pi}) - \cos\{\theta(\boldsymbol{\pi})\}\boldsymbol{y}_0\right], & \theta(\boldsymbol{\pi}) > 0, \\ \boldsymbol{0}, & \theta(\boldsymbol{\pi}) = 0. \end{cases} \tag{S15}$$

Its displayed coordinates are

$$\mathrm{HRIC}_j(\boldsymbol{\pi}) = \frac{\theta(\boldsymbol{\pi})}{\sin\{\theta(\boldsymbol{\pi})\}}\left[\sqrt{\pi_j} - \frac{\cos\{\theta(\boldsymbol{\pi})\}}{\sqrt{p}}\right], \qquad j = 1, \ldots, p, \tag{S16}$$

with the continuous value zero at $\boldsymbol{\pi} = \boldsymbol{\pi}_0$.

**Proposition S2.2** (Riemannian logarithmic map)**.** *For every $\boldsymbol{\pi} \in \overline{\mathcal{S}}^{p-1}$, the vector in Equation* (S15) *is the spherical logarithm* $\mathrm{Log}_{\boldsymbol{y}_0}\{\boldsymbol{y}(\boldsymbol{\pi})\}$.

*Proof.* For $\boldsymbol{y} \neq -\boldsymbol{y}_0$, the spherical logarithm is

$$\mathrm{Log}_{\boldsymbol{y}_0}(\boldsymbol{y}) = \frac{\vartheta}{\sin\vartheta}\left(\boldsymbol{y} - \cos\vartheta\, \boldsymbol{y}_0\right), \qquad \vartheta = \arccos\langle \boldsymbol{y}, \boldsymbol{y}_0\rangle. \tag{S17}$$

Substituting $\boldsymbol{y} = \boldsymbol{y}(\boldsymbol{\pi})$ gives Equation (S15); at $\boldsymbol{\pi} = \boldsymbol{\pi}_0$, both sides equal $\boldsymbol{0}$. □

The map first projects the square-root composition into $\mathcal{H}_p$ and then rescales the projected vector so that its Euclidean norm equals the spherical arc length from the uniform composition.

#### S2.4 Equivariance and contrast structure

**Theorem S2.3** (Scale invariance, permutation equivariance and centring). *Let $\boldsymbol{x} \in \mathbb{R}^p_{\geq 0}$ have positive total, let $b > 0$, and let $R$ be a $p \times p$ permutation matrix. Then*

$$\mathrm{HRIC}\{\mathcal{C}(b\boldsymbol{x})\} = \mathrm{HRIC}\{\mathcal{C}(\boldsymbol{x})\}, \qquad \mathrm{HRIC}(R\boldsymbol{\pi}) = R\,\mathrm{HRIC}(\boldsymbol{\pi}), \tag{S18}$$

*and*

$$\mathbf{1}^{\mathsf{T}}\,\mathrm{HRIC}(\boldsymbol{\pi}) = 0. \tag{S19}$$

*Thus* $\mathrm{HRIC}(\boldsymbol{\pi}) \in \mathcal{H}_p$, *and its effective dimension is* $p-1$.

*Proof.* Scale invariance follows from $\mathcal{C}(b\boldsymbol{x}) = \mathcal{C}(\boldsymbol{x})$. For a permutation, $\boldsymbol{y}(R\boldsymbol{\pi}) = R\boldsymbol{y}(\boldsymbol{\pi})$, $R\boldsymbol{y}_0 = \boldsymbol{y}_0$, and $R$ is orthogonal. The angle and radial scaling in Equation (S15) are therefore unchanged, giving $\mathrm{HRIC}(R\boldsymbol{\pi}) = R\,\mathrm{HRIC}(\boldsymbol{\pi})$. Finally,

$$\sum_{j=1}^{p}\left[\sqrt{\pi_j} - \frac{c(\boldsymbol{\pi})}{\sqrt{p}}\right] = \sum_{j=1}^{p}\sqrt{\pi_j} - \sqrt{p}\,c(\boldsymbol{\pi}) = 0.$$

The common radial scaling preserves this equality. □

Permutation equivariance means that relabelling the features only relabels the coordinates; no feature is privileged as a reference. The $p$ displayed coordinates are linearly dependent through Equation (S19). A full-rank numerical representation can be obtained by multiplying by any $p \times (p-1)$ matrix whose columns form an orthonormal basis of $\mathcal{H}_p$.

#### S2.5 Angular norm, zero compatibility and boundedness

**Theorem S2.4** (Angular norm). *For every* $\boldsymbol{\pi} \in \overline{\mathcal{S}}^{p-1}$,

$$\|\mathrm{HRIC}(\boldsymbol{\pi})\|_2 = \theta(\boldsymbol{\pi}) = \arccos\left\langle\sqrt{\boldsymbol{\pi}}, \sqrt{\boldsymbol{\pi}_0}\right\rangle. \tag{S20}$$

*Proof.* The projected vector has squared norm

$$\|\boldsymbol{y}(\boldsymbol{\pi}) - c(\boldsymbol{\pi})\boldsymbol{y}_0\|_2^2 = 1 - c(\boldsymbol{\pi})^2 = s(\boldsymbol{\pi})^2.$$

Multiplication by $\theta(\boldsymbol{\pi})/\sin\{\theta(\boldsymbol{\pi})\} = \theta(\boldsymbol{\pi})/s(\boldsymbol{\pi})$ gives Equation (S20), including the continuous case $s(\boldsymbol{\pi}) = 0$. □

Define

$$A_p = \arccos\left(p^{-1/2}\right) = \arcsin\sqrt{1 - p^{-1}}. \tag{S21}$$

**Theorem S2.5** (Zero compatibility and global bound). *HRIC is finite and continuous at every point of* $\overline{\mathcal{S}}^{p-1}$, *including all boundary faces, and*

$$0 \leq \|\mathrm{HRIC}(\boldsymbol{\pi})\|_2 \leq A_p < \frac{\pi}{2}. \tag{S22}$$

*The lower bound is attained only at* $\boldsymbol{\pi}_0$, *and the upper bound is attained exactly at the vertices of the simplex.*

*Proof.* Cauchy–Schwarz gives $c(\boldsymbol{\pi}) \leq 1$, with equality only at $\boldsymbol{\pi}_0$. Because $\sqrt{t} \geq t$ on $[0, 1]$, $\sum_{j=1}^{p} \sqrt{\pi_j} \geq 1$, so $c(\boldsymbol{\pi}) \geq p^{-1/2}$. Equality in the latter relation occurs at a simplex vertex. Equation (S20) then gives Equation (S22). Continuity follows from continuity of the square-root map and of the spherical logarithm on the non-negative orthant, which lies within the injectivity radius of $\boldsymbol{y}_0$. □

**Corollary S2.6** (Coordinatewise bound). *For every feature $j$ and every $\boldsymbol{\pi} \in \overline{\mathcal{S}}^{p-1}$,*

$$|\mathrm{HRIC}_j(\boldsymbol{\pi})| \leq \|\mathrm{HRIC}(\boldsymbol{\pi})\|_2 \leq A_p. \tag{S23}$$

*No coordinate diverges when a component vanishes.*

#### S2.6 Invertibility

**Theorem S2.7** (Inverse map and homeomorphism). *HRIC is a homeomorphism from $\overline{\mathcal{S}}^{p-1}$ onto a compact subset of $\mathcal{H}_p$. If $\boldsymbol{z} = \mathrm{HRIC}(\boldsymbol{\pi})$, then $\boldsymbol{\pi} = \boldsymbol{\pi}_0$ when $\boldsymbol{z} = \boldsymbol{0}$. When $\boldsymbol{z} \neq \boldsymbol{0}$, write $\ell = \|\boldsymbol{z}\|_2$ and $\boldsymbol{u} = \boldsymbol{z}/\ell$. The inverse is*

$$\pi_j = \left( \frac{\cos \ell}{\sqrt{p}} + \sin \ell \, u_j \right)^2, \qquad j = 1, \ldots, p. \tag{S24}$$

*The image consists exactly of the vectors $\boldsymbol{z} \in \mathcal{H}_p$ satisfying $0 \leq \|\boldsymbol{z}\|_2 \leq A_p$ for which the expression inside parentheses in Equation* (S24) *is non-negative for every $j$.*

*Proof.* The spherical exponential map reconstructs

$$\boldsymbol{y}(\boldsymbol{\pi}) = \mathrm{Exp}_{\boldsymbol{y}_0}(\boldsymbol{z}) = \cos \ell \, \boldsymbol{y}_0 + \sin \ell \, \boldsymbol{u}.$$

Squaring the coordinates gives Equation (S24). The norm restriction and coordinatewise non-negativity condition are necessary and sufficient for the reconstructed point to lie in the non-negative spherical orthant within the injectivity radius of $\boldsymbol{y}_0$. HRIC is continuous on the compact set $\overline{\mathcal{S}}^{p-1}$, and a continuous bijection from a compact space into the Hausdorff space $\mathcal{H}_p$ is a homeomorphism onto its image. □

Invertibility ensures that the coordinate representation does not discard compositional information. Its image is a bounded domain within the contrast space rather than the full space $\mathcal{H}_p$.

#### S2.7 Local relationship with centred log-ratio coordinates

Define the radial scaling function

$$\lambda(\boldsymbol{\pi}) = \begin{cases} \theta(\boldsymbol{\pi}) / \sin\{\theta(\boldsymbol{\pi})\}, & \theta(\boldsymbol{\pi}) > 0, \\ 1, & \theta(\boldsymbol{\pi}) = 0. \end{cases} \tag{S25}$$

**Proposition S2.8** (Local centred-square-root form). *As $\boldsymbol{\pi} \to \boldsymbol{\pi}_0$,*

$$\lambda(\boldsymbol{\pi}) = 1 + \frac{s(\boldsymbol{\pi})^2}{6} + O\{s(\boldsymbol{\pi})^4\}, \tag{S26}$$

*and*

$$\mathrm{HRIC}(\boldsymbol{\pi}) = [\boldsymbol{y}(\boldsymbol{\pi}) - c(\boldsymbol{\pi}) \boldsymbol{y}_0] \left[ 1 + O\{s(\boldsymbol{\pi})^2\} \right]. \tag{S27}$$

*Proof.* Use the expansion $\arcsin(s) = s + s^3/6 + O(s^5)$ in Equation (S25). $\square$

**Proposition S2.9** (First-order agreement with the centred log-ratio)**.** *Let* $\boldsymbol{\pi} \in \mathcal{S}_+^{p-1}$ *and write* $\pi_j = p^{-1} + \varepsilon_j$*, where* $\sum_{j=1}^p \varepsilon_j = 0$*. As* $\boldsymbol{\varepsilon} \to \mathbf{0}$*,*

$$\mathrm{HRIC}_j(\boldsymbol{\pi}) = \frac{1}{2\sqrt{p}}\,\mathrm{clr}_j(\boldsymbol{\pi}) + o(\|\boldsymbol{\varepsilon}\|_2), \qquad j = 1, \ldots, p, \tag{S28}$$

*where*

$$\mathrm{clr}_j(\boldsymbol{\pi}) = \log \pi_j - \frac{1}{p}\sum_{k=1}^p \log \pi_k. \tag{S29}$$

*Proof.* The expansion $\log(1 + u) = u + o(u)$ gives

$$\mathrm{clr}_j(\boldsymbol{\pi}) = p\varepsilon_j + o(\|\boldsymbol{\varepsilon}\|_2).$$

Similarly,

$$\sqrt{\pi_j} = p^{-1/2} + \frac{\sqrt{p}}{2}\varepsilon_j + o(\|\boldsymbol{\varepsilon}\|_2),$$

and the average linear term vanishes because $\sum_{j=1}^p \varepsilon_j = 0$. Equation (S26) gives $\lambda(\boldsymbol{\pi}) = 1 + o(1)$, so

$$\mathrm{HRIC}_j(\boldsymbol{\pi}) = \frac{\sqrt{p}}{2}\varepsilon_j + o(\|\boldsymbol{\varepsilon}\|_2).$$

Combining the two expansions proves Equation (S28). $\square$

The common positive factor $(2\sqrt{p})^{-1}$ preserves local contrast directions. Statistics invariant to a global coordinate rescaling therefore have the same first-order geometry under HRIC and CLR near the uniform composition. The representations separate towards the boundary, where CLR is undefined and HRIC remains finite.

#### S2.8 Boundary behaviour

**Proposition S2.10** (Finite boundary limits)**.** *Let* $\boldsymbol{\pi}^* \in \overline{\mathcal{S}}^{p-1}$ *be a boundary composition and let* $\mathcal{Z} = \{j : \pi_j^* = 0\}$*. For any sequence* $\boldsymbol{\pi} \to \boldsymbol{\pi}^*$*,*

$$\mathrm{HRIC}_j(\boldsymbol{\pi}) \longrightarrow -\lambda(\boldsymbol{\pi}^*)\frac{c(\boldsymbol{\pi}^*)}{\sqrt{p}} < 0, \qquad j \in \mathcal{Z}. \tag{S30}$$

*By contrast, additive and centred log-ratio coordinates involving a vanishing component diverge to* $-\infty$*.*

*Proof.* For $j \in \mathcal{Z}$, $\sqrt{\pi_j} \to 0$. Substitution into Equation (S16) gives Equation (S30). A log-ratio coordinate contains $\log \pi_j$, which diverges as $\pi_j \to 0^+$. $\square$

At a vertex $\boldsymbol{e}_k$, define

$$\lambda^* = \frac{A_p}{\sqrt{1 - p^{-1}}}. \tag{S31}$$

Then

$$\mathrm{HRIC}_k(\boldsymbol{e}_k) = \lambda^*(1 - p^{-1}), \qquad \mathrm{HRIC}_j(\boldsymbol{e}_k) = -\lambda^* p^{-1} \quad (j \neq k), \tag{S32}$$

and $\|\mathrm{HRIC}(\boldsymbol{e}_k)\|_2 = A_p$.

##### S2.9 Euclidean dissimilarity in the HRIC chart

**Definition S2.11** (HRIC dissimilarity)**.** For $\boldsymbol{\pi}, \boldsymbol{\pi}' \in \overline{\mathcal{S}}^{p-1}$, define

$$d_{\mathrm{HRIC}}(\boldsymbol{\pi}, \boldsymbol{\pi}') = \left\|\mathrm{HRIC}(\boldsymbol{\pi}) - \mathrm{HRIC}(\boldsymbol{\pi}')\right\|_2. \tag{S33}$$

**Proposition S2.12** (Metric properties and bounds)**.** *The function $d_{\mathrm{HRIC}}$ is a metric on $\overline{\mathcal{S}}^{p-1}$,*

$$0 \le d_{\mathrm{HRIC}}(\boldsymbol{\pi}, \boldsymbol{\pi}') \le 2A_p, \tag{S34}$$

*and*

$$d_{\mathrm{HRIC}}(\boldsymbol{\pi}, \boldsymbol{\pi}_0) = \|\mathrm{HRIC}(\boldsymbol{\pi})\|_2 = \theta(\boldsymbol{\pi}, \boldsymbol{\pi}_0). \tag{S35}$$

*Proof.* Non-negativity, symmetry and the triangle inequality follow from Euclidean distance. Identity of indiscernibles follows from Theorem S2.7, and the global bound follows from the triangle inequality and Theorem S2.5. Equation (S35) follows from $\mathrm{HRIC}(\boldsymbol{\pi}_0) = \mathbf{0}$ and Theorem S2.4. □

HRIC is an exact radial isometry about the uniform composition, not a globally distance-preserving flattening of the sphere. For two general points, let

$$\ell = \|\mathrm{HRIC}(\boldsymbol{\pi})\|_2, \qquad \ell' = \left\|\mathrm{HRIC}(\boldsymbol{\pi}')\right\|_2,$$

and let $\psi$ be the angle between their HRIC vectors. Then

$$d_{\mathrm{HRIC}}(\boldsymbol{\pi}, \boldsymbol{\pi}')^2 = \ell^2 + (\ell')^2 - 2\ell\ell' \cos\psi. \tag{S36}$$

Normal coordinates preserve the Riemannian metric at the base point, so pairwise spherical distances among compositions close to $\boldsymbol{\pi}_0$ agree with $d_{\mathrm{HRIC}}$ to first order. Equation (S28) also gives

$$\mathrm{HRIC}(\boldsymbol{\pi}) - \mathrm{HRIC}(\boldsymbol{\pi}') = \frac{1}{2\sqrt{p}}\left[\mathrm{clr}(\boldsymbol{\pi}) - \mathrm{clr}(\boldsymbol{\pi}')\right] + o\left\{\|\boldsymbol{\pi} - \boldsymbol{\pi}_0\|_2 + \left\|\boldsymbol{\pi}' - \boldsymbol{\pi}_0\right\|_2\right\}. \tag{S37}$$

##### S2.10 Reference-based coordinates and relation to log ratios

**Definition S2.13** (Reference-based HRIC)**.** Choose a reference component $r \in \{1, \dots, p\}$. For $j \ne r$, define

$$\mathrm{RHRIC}_j^{(r)}(\boldsymbol{\pi}) = \lambda(\boldsymbol{\pi})\left(\sqrt{\pi_j} - \sqrt{\pi_r}\right). \tag{S38}$$

The reference-based representation has $p-1$ displayed coordinates and remains finite if $\pi_j = 0$ or $\pi_r = 0$. It is scale invariant and bounded because

$$\left|\mathrm{RHRIC}_j^{(r)}(\boldsymbol{\pi})\right| \le \lambda(\boldsymbol{\pi})\left|\sqrt{\pi_j} - \sqrt{\pi_r}\right| \le \frac{\pi}{2}. \tag{S39}$$

It depends on the selected reference and is not permutation symmetric in the same sense as centred HRIC. The centred and reference-based coordinates encode the same scaled square-root contrasts:

$$\mathrm{HRIC}_j(\boldsymbol{\pi}) - \mathrm{HRIC}_k(\boldsymbol{\pi}) = \lambda(\boldsymbol{\pi})\left(\sqrt{\pi_j} - \sqrt{\pi_k}\right), \tag{S40}$$

and, for $j \ne r$,

$$\mathrm{RHRIC}_j^{(r)}(\boldsymbol{\pi}) = \mathrm{HRIC}_j(\boldsymbol{\pi}) - \mathrm{HRIC}_r(\boldsymbol{\pi}). \tag{S41}$$

The relation between HRIC and RHRIC parallels that between CLR and ALR: one is centred and reference-free, whereas the other records the same pairwise contrasts relative to a chosen component.

Near the uniform composition,

$$\mathrm{RHRIC}_j^{(r)}(\boldsymbol{\pi}) = \frac{1}{2\sqrt{p}}\left(\log\pi_j - \log\pi_r\right) + o\left(\|\boldsymbol{\pi} - \boldsymbol{\pi}_0\|_2\right), \qquad j \ne r. \tag{S42}$$

Thus RHRIC and ALR share the same local contrast direction up to the global factor $(2\sqrt{p})^{-1}$.

##### S2.11 Complementarity with Aitchison geometry

Aitchison geometry is the canonical log-ratio geometry of the open simplex [17, 18]. It provides perturbation and powering operations, subcompositional coherence and orthonormal ILR coordinates. HRIC serves a different role: it supplies a finite, bounded and permutation-symmetric normal-coordinate chart on the closed simplex, where log-ratio coordinates cannot be evaluated without replacing zeros.

HRIC is not generally subcompositionally coherent. Removing features and re-closing changes $p$, $\boldsymbol{y}_0$, $c(\boldsymbol{\pi})$ and $A_p$, so the coordinates of retained parts need not equal their coordinates in the full composition. The intended use is therefore a fixed feature dictionary shared by all communities in a comparison.

Supplementary Table 1: Selected properties of compositional coordinate systems. Effective dimension counts linearly independent coordinates. Symmetry indicates whether relabelling parts only relabels the displayed coordinates without selecting a privileged reference or basis.

| Method | Domain | Effective dimension | Exact zeros | Bounded | Symmetric | Reference or basis |
|---|---|---|---|---|---|---|
| CLR | Open simplex | $p-1$ | No | No | Yes | None |
| ILR | Open simplex | $p-1$ | No | No | Basis-dependent | Orthonormal basis |
| ALR | Open simplex | $p-1$ | No | No | No | Reference part |
| HRIC | Closed simplex | $p-1$ | Yes | Yes | Yes | Uniform direction |
| RHRIC | Closed simplex | $p-1$ | Yes | Yes | No | Reference part |

#### Supplementary Note 3: Diversity partitioning, pairwise dissimilarity, predictor-associated HRIC dispersion and simulation evaluation

##### S3.1 Simplex Hellinger alpha diversity

**Definition S3.1** (Simplex Hellinger alpha diversity)**.** For a composition $\boldsymbol{\pi} \in \overline{\mathcal{S}}^{p-1}$, define

$$\alpha_{\mathrm{SH}}(\boldsymbol{\pi}) = 1 - \frac{\|\mathrm{HRIC}(\boldsymbol{\pi})\|_2^2}{A_p^2}. \tag{S43}$$

By Theorem S2.5, $\alpha_{\mathrm{SH}}(\boldsymbol{\pi}) \in [0, 1]$. It equals 1 only at the uniform composition and 0 exactly at a simplex vertex. It is therefore a normalized angular evenness index: lower values indicate greater concentration of compositional mass. Its complement decomposes additively by feature,

$$1 - \alpha_{\mathrm{SH}}(\boldsymbol{\pi}) = \sum_{j=1}^{p} \frac{\mathrm{HRIC}_j(\boldsymbol{\pi})^2}{A_p^2}. \tag{S44}$$

Each term is a taxon's contribution to squared angular departure from uniformity, not its relative abundance and not a feature-wise causal effect.

**Proposition S3.2** (Relation to the Hill number of order one half)**.** *Let*

$${}^{1/2}D(\boldsymbol{\pi}) = \left( \sum_{j=1}^{p} \sqrt{\pi_j} \right)^2 \tag{S45}$$

*be the Hill number of order* $1/2$ *[3]. Then*

$$\alpha_{\mathrm{SH}}(\boldsymbol{\pi}) = 1 - \frac{\arccos^2 \sqrt{{}^{1/2}D(\boldsymbol{\pi})/p}}{A_p^2}. \tag{S46}$$

*Consequently,* $\alpha_{\mathrm{SH}}$ *is a strictly increasing bounded reparameterization of* ${}^{1/2}D$*.*

*Proof.* Equation (S14) gives

$$c(\boldsymbol{\pi}) = p^{-1/2} \sum_{j=1}^{p} \sqrt{\pi_j} = \sqrt{{}^{1/2}D(\boldsymbol{\pi})/p}.$$

Apply Theorem S2.4 in Definition S3.1. Because $\arccos(c)$ decreases in $c$, alpha diversity increases with the Hill number. □

#### S3.2 Pooled regional composition and gamma diversity

Let

$$\boldsymbol{\Pi} = (\boldsymbol{\pi}_1, \ldots, \boldsymbol{\pi}_n)^{\mathsf{T}}$$

denote a regional ensemble of local compositions, and let the weights $w_1, \ldots, w_n$ sum to one. The pooled regional composition $\boldsymbol{\pi}_\gamma$ is defined by Equation (S7). The package default uses equal weights $w_i = 1/n$, so every local community contributes equally regardless of sequencing depth.

**Definition S3.3** (Simplex Hellinger gamma diversity)**.** The diversity of the pooled regional composition is

$$\gamma_{\mathrm{SH}}(\boldsymbol{\Pi}; \boldsymbol{w}) = 1 - \frac{\|\mathrm{HRIC}(\boldsymbol{\pi}_\gamma)\|_2^2}{A_p^2}. \tag{S47}$$

For equal weights, write $\gamma_{\mathrm{SH}}(\boldsymbol{\Pi})$.

The pooled regional composition differs from the composition obtained by pooling all reads, which would weight samples in proportion to sequencing depth. Alternative ecological sampling weights can be supplied, but they define a different estimand from the equal-community-weight default.

#### S3.3 Additive regional beta diversity

**Definition S3.4** (Simplex Hellinger regional beta diversity)**.** The weighted additive regional beta diversity is

$$\beta_{\mathrm{SH}}(\boldsymbol{\Pi}; \boldsymbol{w}) = \gamma_{\mathrm{SH}}(\boldsymbol{\Pi}; \boldsymbol{w}) - \sum_{i=1}^{n} w_i \alpha_{\mathrm{SH}}(\boldsymbol{\pi}_i). \tag{S48}$$

For equal weights,

$$\beta_{\mathrm{SH}}(\boldsymbol{\Pi}) = \gamma_{\mathrm{SH}}(\boldsymbol{\Pi}) - \frac{1}{n} \sum_{i=1}^{n} \alpha_{\mathrm{SH}}(\boldsymbol{\pi}_i). \tag{S49}$$

The regional beta component measures the increase in normalized diversity produced by pooling the local compositions into the regional ensemble. For equal weights, the additive identity is

$$\gamma_{\mathrm{SH}}(\boldsymbol{\Pi}) = \frac{1}{n} \sum_{i=1}^{n} \alpha_{\mathrm{SH}}(\boldsymbol{\pi}_i) + \beta_{\mathrm{SH}}(\boldsymbol{\Pi}). \tag{S50}$$

**Theorem S3.5** (Non-negativity of the additive regional beta component)**.** *For every regional ensemble,*

$$0 \le \beta_{\mathrm{SH}}(\boldsymbol{\Pi}; \boldsymbol{w}) \le 1. \tag{S51}$$

*If all weights are positive,* $\beta_{\mathrm{SH}}(\boldsymbol{\Pi}; \boldsymbol{w}) = 0$ *if and only if all included local compositions are identical.*

*Proof.* Write

$$c_i = \boldsymbol{y}_i^{\mathsf{T}} \boldsymbol{y}_0, \qquad c_\gamma = \boldsymbol{y}_\gamma^{\mathsf{T}} \boldsymbol{y}_0.$$

From Equation (S7),

$$c_\gamma = \frac{\sum_{i=1}^n w_i c_i}{\|\bar{\boldsymbol{y}}_w\|_2} \ge \sum_{i=1}^n w_i c_i, \tag{S52}$$

because $\|\bar{\boldsymbol{y}}_w\|_2 \le \sum_{i=1}^n w_i \|\boldsymbol{y}_i\|_2 = 1$. Let $f(c) = \arccos^2(c)$ on $[0,1]$. The function is decreasing and convex because, writing $c = \cos t$,

$$f''(c) = \frac{2\left\{\sqrt{1-c^2} - c \arccos(c)\right\}}{(1-c^2)^{3/2}} = \frac{2(\sin t - t\cos t)}{\sin^3 t} \ge 0. \tag{S53}$$

Therefore,

$$\|\mathrm{HRIC}(\boldsymbol{\pi}_\gamma)\|_2^2 = f(c_\gamma) \le f\left(\sum_{i=1}^n w_i c_i\right) \le \sum_{i=1}^n w_i f(c_i) = \sum_{i=1}^n w_i \|\mathrm{HRIC}(\boldsymbol{\pi}_i)\|_2^2 . \tag{S54}$$

Substitution into Equations (S43) and (S48) proves non-negativity. The upper bound follows because every squared norm is at most $A_p^2$. Equality at zero requires $\|\bar{\boldsymbol{y}}_w\|_2 = 1$, which, for positive weights and unit vectors, occurs exactly when all $\boldsymbol{y}_i$, and hence all $\boldsymbol{\pi}_i$, are identical. □

**Remark S3.6** (Regional beta diversity is not coordinate variance)**.** In general,

$$\beta_{\mathrm{SH}}(\boldsymbol{\Pi}) \ne \frac{1}{nA_p^2} \sum_{i=1}^n \|\boldsymbol{z}_i - \bar{\boldsymbol{z}}\|_2^2 , \qquad \boldsymbol{z}_i = \mathrm{HRIC}(\boldsymbol{\pi}_i), \qquad \bar{\boldsymbol{z}} = \frac{1}{n}\sum_{i=1}^n \boldsymbol{z}_i. \tag{S55}$$

The left-hand side compares mean local angular diversity with diversity of the pooled regional composition. The right-hand side is Euclidean dispersion around the arithmetic mean of the HRIC coordinates. Both are meaningful but answer different ecological questions.

**Remark S3.7** (Signed local–regional diversity gap)**.** For sample $i$, define

$$\tau_i = \gamma_{\mathrm{SH}}(\boldsymbol{\Pi}) - \alpha_{\mathrm{SH}}(\boldsymbol{\pi}_i) = \frac{\|\boldsymbol{z}_i\|_2^2 - \|\mathrm{HRIC}(\boldsymbol{\pi}_\gamma)\|_2^2}{A_p^2}. \tag{S56}$$

Its regional mean is

$$\frac{1}{n}\sum_{i=1}^n \tau_i = \beta_{\mathrm{SH}}(\boldsymbol{\Pi}).$$

Individual values can be negative when a local community is more even than the pooled regional composition. The quantity is therefore a signed local–regional diversity gap rather than a non-negative distance or an independent measure of taxonomic replacement. A non-negative sample-to-region pairwise dissimilarity is $d_{\mathrm{HRIC}}(\boldsymbol{\pi}_i, \boldsymbol{\pi}_\gamma)$.

##### S3.4 Pairwise dissimilarity and predictor-associated HRIC dispersion

The HRIC dissimilarity in Equation (S33) is defined for a pair of local communities. By contrast, $\beta_{\mathrm{SH}}$ is one scalar for the complete regional ensemble. A matrix of pairwise HRIC dissimilarities can visualize or model compositional variation, but no individual entry of that matrix is the regional beta component.

For a categorical condition $g_i \in \{1, \ldots, G\}$, let

$$W_h = \sum_{i:g_i=h} w_i,$$

define the condition-specific centroid

$$\bar{\boldsymbol{z}}_h = W_h^{-1} \sum_{i:g_i=h} w_i \boldsymbol{z}_i, \tag{S57}$$

and define the overall centroid

$$\bar{\boldsymbol{z}} = \sum_{i=1}^{n} w_i \boldsymbol{z}_i.$$

**Definition S3.8** (Condition-associated HRIC dispersion)**.** The between-condition component is

$$\delta_{\mathrm{SH}}(\boldsymbol{\Pi}, \boldsymbol{g}; \boldsymbol{w}) = \sum_{h=1}^{G} W_h \left\| \bar{\boldsymbol{z}}_h - \bar{\boldsymbol{z}} \right\|_2^2 . \tag{S58}$$

For equal weights, the same quantity can be written as

$$\delta_{\mathrm{SH}}(\boldsymbol{\Pi}, \boldsymbol{g}) = V_{\mathrm{region}} - \sum_{h=1}^{G} \frac{n_h}{n} V_h, \tag{S59}$$

where

$$V_{\mathrm{region}} = \frac{1}{n} \sum_{i=1}^{n} \left\| \boldsymbol{z}_i - \bar{\boldsymbol{z}} \right\|_2^2 , \qquad V_h = \frac{1}{n_h} \sum_{i:g_i=h} \left\| \boldsymbol{z}_i - \bar{\boldsymbol{z}}_h \right\|_2^2 . \tag{S60}$$

**Proposition S3.9** (Euclidean analysis-of-variance identity)**.** *For equal weights,*

$$SS_{\mathrm{between}} = \sum_{h=1}^{G} n_h \left\| \bar{\boldsymbol{z}}_h - \bar{\boldsymbol{z}} \right\|_2^2 = n\, \delta_{\mathrm{SH}}(\boldsymbol{\Pi}, \boldsymbol{g}), \tag{S61}$$

*and*

$$SS_{\mathrm{total}} = SS_{\mathrm{within}} + SS_{\mathrm{between}}. \tag{S62}$$

*Proof.* For sample $i$ in condition $h$, write

$$\boldsymbol{z}_i - \bar{\boldsymbol{z}} = (\boldsymbol{z}_i - \bar{\boldsymbol{z}}_h) + (\bar{\boldsymbol{z}}_h - \bar{\boldsymbol{z}}).$$

Squaring and summing within each condition eliminates the cross term because $\sum_{i:g_i=h}(\boldsymbol{z}_i - \bar{\boldsymbol{z}}_h) = \boldsymbol{0}$. Summing over conditions gives Equations (S59)–(S62) and establishes non-negativity. □

A dimensionless normalized effect size is

$$\delta_{\mathrm{SH}}^{*}(\mathbf{\Pi}, \boldsymbol{g}) = \frac{\delta_{\mathrm{SH}}(\mathbf{\Pi}, \boldsymbol{g})}{A_p^2}, \tag{S63}$$

so

$$SS_{\mathrm{between}} = nA_p^2 \delta_{\mathrm{SH}}^{*}. \tag{S64}$$

This effect size measures separation of condition-specific HRIC centroids. It is distinct from regional beta diversity in Equation (S49), which compares mean local diversity with diversity of the pooled regional composition.

The same construction extends to continuous predictors and covariate adjustment. Let $\boldsymbol{Z}$ be the $n \times p$ HRIC matrix, let $M_0$ contain an intercept and nuisance covariates, and let $M_1 = [M_0 \; L]$ additionally contain the focal environmental or host variables. If $P_0$ and $P_1$ are the orthogonal projection matrices onto the column spaces of $M_0$ and $M_1$, respectively, the additional model-explained HRIC dispersion is

$$\delta_{\mathrm{SH}}(L \mid M_0) = \frac{1}{n} \operatorname{tr}\left\{ \boldsymbol{Z}^{\mathsf{T}} (P_1 - P_0) \boldsymbol{Z} \right\}. \tag{S65}$$

For nested full-rank column spaces, $P_1 - P_0$ is positive semidefinite, so Equation (S65) is non-negative. Continuous gradients, multi-level factors and interactions can therefore be represented in the same coordinate geometry.

#### S3.5 Connection with PERMANOVA

Let $\mathbf{\Delta}^{(2)}$ contain squared HRIC dissimilarities and let

$$C_n = I_n - n^{-1}\mathbf{1}\mathbf{1}^{\mathsf{T}}.$$

The Gower matrix is

$$\boldsymbol{B} = -\frac{1}{2} C_n \mathbf{\Delta}^{(2)} C_n. \tag{S66}$$

Because HRIC dissimilarities are Euclidean,

$$\boldsymbol{B} = \boldsymbol{Z}_c \boldsymbol{Z}_c^{\mathsf{T}}, \qquad \boldsymbol{Z}_c = C_n \boldsymbol{Z}. \tag{S67}$$

For a group design with full-model projection $P_1$ and intercept-only projection $P_0$, the PERMANOVA numerator is

$$\operatorname{tr}\{(P_1 - P_0)\boldsymbol{B}\} = \operatorname{tr}\left\{ \boldsymbol{Z}^{\mathsf{T}} (P_1 - P_0) \boldsymbol{Z} \right\} = SS_{\mathrm{between}} = n\,\delta_{\mathrm{SH}}. \tag{S68}$$

PERMANOVA divides this numerator, after degrees-of-freedom scaling, by residual within-model dispersion and obtains a permutation $P$ value [13, 14]. Thus, $\delta_{\mathrm{SH}}$ is the condition-associated HRIC-dispersion effect size, whereas PERMANOVA tests the same numerator relative to residual heterogeneity. Within-condition multivariate spread should be examined separately because permutation results can be sensitive to differences in both centroid location and within-condition spread [10].

Supplementary Table 2: Distinct ecological and geometric quantities in the HRIC framework.

| Quantity | Unit of analysis | Interpretation |
|---|---|---|
| $\alpha_{\mathrm{SH}}(\boldsymbol{\pi}_i)$ | One local community | Within-community evenness, normalized from squared angular departure from uniformity. |
| $\gamma_{\mathrm{SH}}(\boldsymbol{\Pi})$ | One regional ensemble | Diversity of the equal-community-weighted pooled regional composition. |
| $\beta_{\mathrm{SH}}(\boldsymbol{\Pi})$ | One regional ensemble | Additive local–regional differentiation, $\gamma - \bar{\alpha}$. |
| $\tau_i$ | One local community within a region | Signed local–regional diversity gap, $\gamma - \alpha_i$. |
| $d_{\mathrm{HRIC}}(\boldsymbol{\pi}_i, \boldsymbol{\pi}_j)$ | One pair of communities | Euclidean dissimilarity between HRIC coordinate vectors. |
| $\delta_{\mathrm{SH}}(\boldsymbol{\Pi}, \boldsymbol{g})$ | One predictor in a region | Between-centroid or model-explained HRIC dispersion associated with the predictor. |
| PERMANOVA $P$ value | One model term | Permutation evidence that predictor-explained dissimilarity variation is large relative to residual variation. |

#### S3.6 Practical interpretation and scope

The framework follows a fixed order of ecological questions:

1. define the regional ensemble and hold its feature dictionary fixed;
2. use $\alpha_{\mathrm{SH}}$ to summarize each local community;
3. use $\gamma_{\mathrm{SH}}$ and $\beta_{\mathrm{SH}}$ to summarize the complete local–regional hierarchy;
4. use $d_{\mathrm{HRIC}}$ to describe pairwise compositional dissimilarity; and
5. use $\delta_{\mathrm{SH}}$, Equation (S65) or another model-based sum of squares to quantify the component associated with a focal predictor, followed by PERMANOVA or another valid inferential procedure.

HRIC is a normal-coordinate chart centred at the uniform composition rather than a globally distance-preserving flattening of the sphere. Pairwise Euclidean dissimilarity is exact along radii from uniform and locally faithful for general pairs. HRIC is not subcompositionally coherent and should not be compared across changing feature dictionaries without redefining the regional ensemble. Observed zeros are admitted geometrically, but HRIC alone does not determine whether a zero represents structural absence or finite-sampling non-detection. Predictor-associated multivariate evidence also does not establish causality without an appropriate design and adequate control of confounding.

#### S3.7 Simulation template and perturbation mechanisms

We fitted SparseDOSSA2 (v.0.99.2) to the throat 16S rRNA gene OTU table and phylogeny distributed with the `GUniFrac` package [19–21]. Taxa occurring in fewer than two reference samples were removed, leaving 616 taxa. Patristic distances were calculated from the pruned phylogeny, and balanced polar-ordination clustering was used to construct phylogenetically coherent and approximately size-balanced taxon groups.

Each simulated dataset contained 100 samples. Binary condition labels were drawn independently from a Bernoulli distribution with probability 0.5. For every taxon in a selected cluster, a positive SparseDOSSA2 effect shifted the fitted log-scale abundance mean, the log odds of taxon presence or both. These settings generated abundance-only, prevalence-only and joint abundance–prevalence perturbations. The effect parameter ranged from 0 to 1 in increments of 0.05.

##### S3.8 Geometric correspondence analysis

For the correspondence analysis, the reference phylogeny was divided into 12 taxon clusters, and each cluster was used in turn as the perturbed set. For each of the three perturbation modes and each of the 21 effect sizes, 125 independent simulation replicates were generated. Within each replicate, the 12 clusters were perturbed in separate datasets, and the resulting quantities were averaged across clusters.

Normalized condition-associated HRIC dispersion in Supplementary Fig. 1 is the dimensionless quantity in Equation (S63). The between-condition and within-condition sums of squares were calculated directly from HRIC coordinates. The exact identity in Equation (S64) was compared with the trace-based pseudo-$F$ statistic

$$F_{\mathrm{HRIC}} = \frac{SS_{\mathrm{between}}/(G-1)}{SS_{\mathrm{within}}/(n-G)}. \tag{S69}$$

For $n = 100$ samples and $p = 616$ taxa, the theoretical scale factor was

$$nA_p^2 = 234.2413, \tag{S70}$$

reported as 234.2 in Supplementary Fig. 1. Across all correspondence simulations, the largest absolute discrepancy from Equation (S64) was $6.1 \times 10^{-14}$. The numerical implementation therefore matched the analytic identity to machine precision. The relationships between normalized condition-associated HRIC dispersion and the complete pseudo-$F$ statistic had $R^2 = 0.998$, 0.999 and 0.993 under abundance, joint and prevalence perturbations, respectively. The relationship with pseudo-$F$ is not an identity because pseudo-$F$ additionally incorporates within-condition heterogeneity.

##### S3.9 Power analysis

For the power analysis, the reference phylogeny was divided into $k = 2, 4, 6$ or 8 clusters, and every cluster was used in turn as the perturbed set. For each combination of phylogenetic partition, perturbed cluster, perturbation mode and effect size, 100 independent datasets containing 100 samples were generated. Supplementary Fig. 1c displays the cluster containing the largest number of taxa for each value of $k$; size ties were resolved using fitted abundance rank. The displayed clusters contained 308, 154, 103 and 77 taxa for $k = 2, 4, 6$ and 8, respectively.

HRIC coordinates were calculated directly with `HRIC::HRIC` (v.0.1.0), and pairwise Euclidean dissimilarities between coordinates were used without zero replacement. Comparator dissimilarities were Bray–Curtis, ordinary Euclidean and binary Jaccard, together with Aitchison and ilr–Euclidean dissimilarities after adding pseudocounts of 0.01, 0.1, 0.5 or 1. Weighted, unweighted and generalized UniFrac dissimilarities were also evaluated but are not displayed in Supplementary Fig. 1.

For a fixed pseudocount, Euclidean distance in an orthonormal ILR basis is geometrically identical to Aitchison distance. The paired implementations were retained as a computational check. Small differences in empirical rejection proportions arose from independent permutation streams rather than different underlying geometries.

Community-level association was tested with `vegan::adonis2` (v.2.7-5) using 999 permutations [14]. Empirical power was the proportion of replicate permutation $P$ values below 0.05 at non-zero effect sizes. For visualization, each empirical power series was fitted with a binomial generalized additive model using a shrinkage cubic regression spline, and the displayed fit was constrained to be non-decreasing with effect size.

Under abundance-only perturbations, HRIC dissimilarity reached 80% empirical power at effect sizes of 0.75, 0.75, 0.75 and 0.80 for $k = 2, 4, 6$ and 8, respectively. The corresponding thresholds for Bray–Curtis were 0.70, 0.75, 0.75 and 0.70. None of the displayed Aitchison, ILR–Euclidean or binary Jaccard variants reached 80% power over the simulated effect range.

Under joint abundance–prevalence perturbations, HRIC dissimilarity reached 80% power at effect sizes of 0.50–0.55, while several log-ratio and occurrence-sensitive dissimilarities reached the same threshold at similar or smaller effects. Under prevalence-only perturbations at an effect size of 0.50, power for HRIC dissimilarity ranged from 0.07 to 0.11 across the four phylogenetic partitions, compared with 0.48–1.00 for binary Jaccard and 0.39–0.79 for Aitchison dissimilarity after adding a pseudocount of 0.5.

##### S3.10 Null calibration

Type I error was the proportion of permutation $P$ values below 0.05 at effect size zero. To match Supplementary Fig. 1c, the largest taxon cluster was selected within each combination of $k$ and nominal perturbation mode, with fitted abundance rank used to resolve size ties. Each null setting contained 100 independently simulated datasets.

The mean type I error of PERMANOVA based on HRIC dissimilarities was 0.0533 across the 12 combinations of perturbation mode and phylogenetic partition, with a setting-specific range of 0.0200–0.0900. Across all evaluated methods, overall mean type I error ranged from 0.0367 to 0.0600. With 100 null replicates per cell, the Monte Carlo standard error at a true rejection probability of 0.05 is approximately 0.022; setting-specific ranges should therefore be interpreted together with method-level averages.

##### S3.11 Simulation results

Regional beta diversity and condition-associated HRIC dispersion answer complementary ecological questions. Simplex Hellinger regional beta diversity summarizes local–regional differentiation for all communities in a prespecified region. For a categorical condition, `HRIC::SHdelta` instead quantifies condition-associated HRIC dispersion, defined by the weighted displacement of condition-specific HRIC centroids from the overall centroid. PERMANOVA based on HRIC dissimilarities evaluates the same between-condition sum of squares relative to the compositional heterogeneity that remains within conditions [14]. Thus, `SHdelta` is the effect-size counterpart of the PERMANOVA numerator, whereas PERMANOVA supplies the permutation test. The geometric derivation is given in Supplementary Note 3.

We examined this correspondence and its sensitivity to different modes of ecological change using zero-inflated microbial-community simulations grounded in an empirical throat microbiome. SparseDOSSA2 was fitted to the 16S rRNA gene data and phylogeny distributed with the `GUniFrac` package, leaving 616 taxa after filtering [19–21]. The phylogeny was divided into coherent taxon clusters, and one cluster at a time was perturbed through changes in abundance, prevalence or both. Abundance perturbations redistributed compositional mass among taxa, prevalence perturbations

Supplementary Table 3: Empirical type I error of dissimilarity-based PERMANOVA in null simulations. Values under each nominal perturbation mode are mean rejection proportions across $k = 2, 4, 6$ and 8 at effect size zero. The final column reports the overall mean and range across all 12 combinations of perturbation mode and $k$. Each combination contained 100 independently simulated datasets, and tests used a nominal level of 0.05. Parenthetical values denote the pseudocount added before log-ratio transformation.

| Method | Abundance | Joint | Prevalence | Overall mean (range) |
|---|---|---|---|---|
| HRIC dissimilarity | 0.0525 | 0.0500 | 0.0575 | 0.0533 (0.0200–0.0900) |
| Bray–Curtis | 0.0600 | 0.0450 | 0.0500 | 0.0517 (0.0200–0.1000) |
| Euclidean | 0.0450 | 0.0300 | 0.0350 | 0.0367 (0.0100–0.0700) |
| Jaccard | 0.0375 | 0.0425 | 0.0700 | 0.0500 (0.0100–0.0800) |
| Aitchison (0.01) | 0.0425 | 0.0550 | 0.0675 | 0.0550 (0.0200–0.1000) |
| Aitchison (0.1) | 0.0500 | 0.0600 | 0.0625 | 0.0575 (0.0100–0.1000) |
| Aitchison (0.5) | 0.0500 | 0.0600 | 0.0600 | 0.0567 (0.0200–0.1000) |
| Aitchison (1) | 0.0525 | 0.0675 | 0.0500 | 0.0567 (0.0200–0.1000) |
| ilr–Euclidean (0.01) | 0.0475 | 0.0600 | 0.0625 | 0.0567 (0.0200–0.1000) |
| ilr–Euclidean (0.1) | 0.0525 | 0.0550 | 0.0650 | 0.0575 (0.0200–0.1100) |
| ilr–Euclidean (0.5) | 0.0550 | 0.0625 | 0.0625 | 0.0600 (0.0200–0.1200) |
| ilr–Euclidean (1) | 0.0500 | 0.0600 | 0.0500 | 0.0533 (0.0200–0.1000) |

changed how frequently taxa occurred across communities, and joint perturbations combined both processes.

We first verified the geometric correspondence using 12 phylogenetic clusters and 100 samples per simulated dataset. Each cluster was perturbed in turn, and the resulting quantities were averaged across clusters within each replicate. The numerical results matched the exact proportionality between normalized condition-associated HRIC dispersion and the PERMANOVA between-condition sum of squares: for 616 taxa and 100 samples, the proportionality constant was 234.2, and the largest absolute numerical error was $6.1 \times 10^{-14}$ (Supplementary Fig. 1b). Normalized condition-associated HRIC dispersion also tracked the complete PERMANOVA pseudo-$F$ statistic closely under abundance, joint and prevalence perturbations ($R^2 = 0.998$, 0.999 and 0.993, respectively; Supplementary Fig. 1a). The small departure from exact proportionality with pseudo-$F$ arises because pseudo-$F$ additionally incorporates within-condition heterogeneity. Condition-associated HRIC dispersion therefore records the magnitude of centroid separation, whereas pseudo-$F$ places that separation in the context of residual community variation.

We next evaluated how this shared geometry responds to different simulated signals. The reference phylogeny was divided into 2, 4, 6 or 8 approximately size-balanced clusters, and the largest cluster at each partition resolution was used for the comparison shown in Supplementary Fig. 1c. Under abundance-only perturbations, HRIC dissimilarity closely tracked Bray–Curtis, reaching 80% empirical power at effect sizes of 0.75, 0.75, 0.75 and 0.80 for 2, 4, 6 and 8 clusters, respectively. The corresponding Bray–Curtis thresholds were 0.70, 0.75, 0.75 and 0.70, whereas none of the displayed Aitchison, ilr–Euclidean or Jaccard variants reached 80% power. Under joint abundance–prevalence perturbations, HRIC dissimilarity reached 80% power at effect sizes of 0.50–0.55, with several log-ratio and occurrence-sensitive dissimilarities detecting changes at similar or smaller effect sizes.

Prevalence-only perturbations produced a different ordering. At an effect size of 0.50, power for HRIC dissimilarity was 0.07–0.11 across the four phylogenetic partitions, compared with 0.48–1.00 for presence–absence Jaccard and 0.39–0.79 for Aitchison dissimilarity with a pseudocount of 0.5.

These patterns distinguish the ecological information emphasized by the different geometries. HRIC dissimilarity primarily captures abundance-weighted redistribution of compositional mass, including abundance changes and joint abundance–occurrence changes, whereas binary Jaccard directly targets differences in community membership. Their contrasting performance under prevalence-only perturbation therefore reflects different ecological targets rather than different tests of an identical quantity.

PERMANOVA based on HRIC dissimilarities remained close to its nominal level under the global null. Its mean type I error was 0.0533 across the 12 combinations of perturbation mode and phylogenetic partition, with a setting-specific range of 0.0200–0.0900. Across all methods shown in Supplementary Fig. 1c, overall mean type I error ranged from 0.0367 to 0.0600 (Supplementary Table 3). Together, these simulations verify the computational correspondence between condition-associated HRIC dispersion and the between-condition component of PERMANOVA and show how PERMANOVA based on HRIC dissimilarities distinguishes abundance-weighted compositional redistribution from occurrence-based community differences.

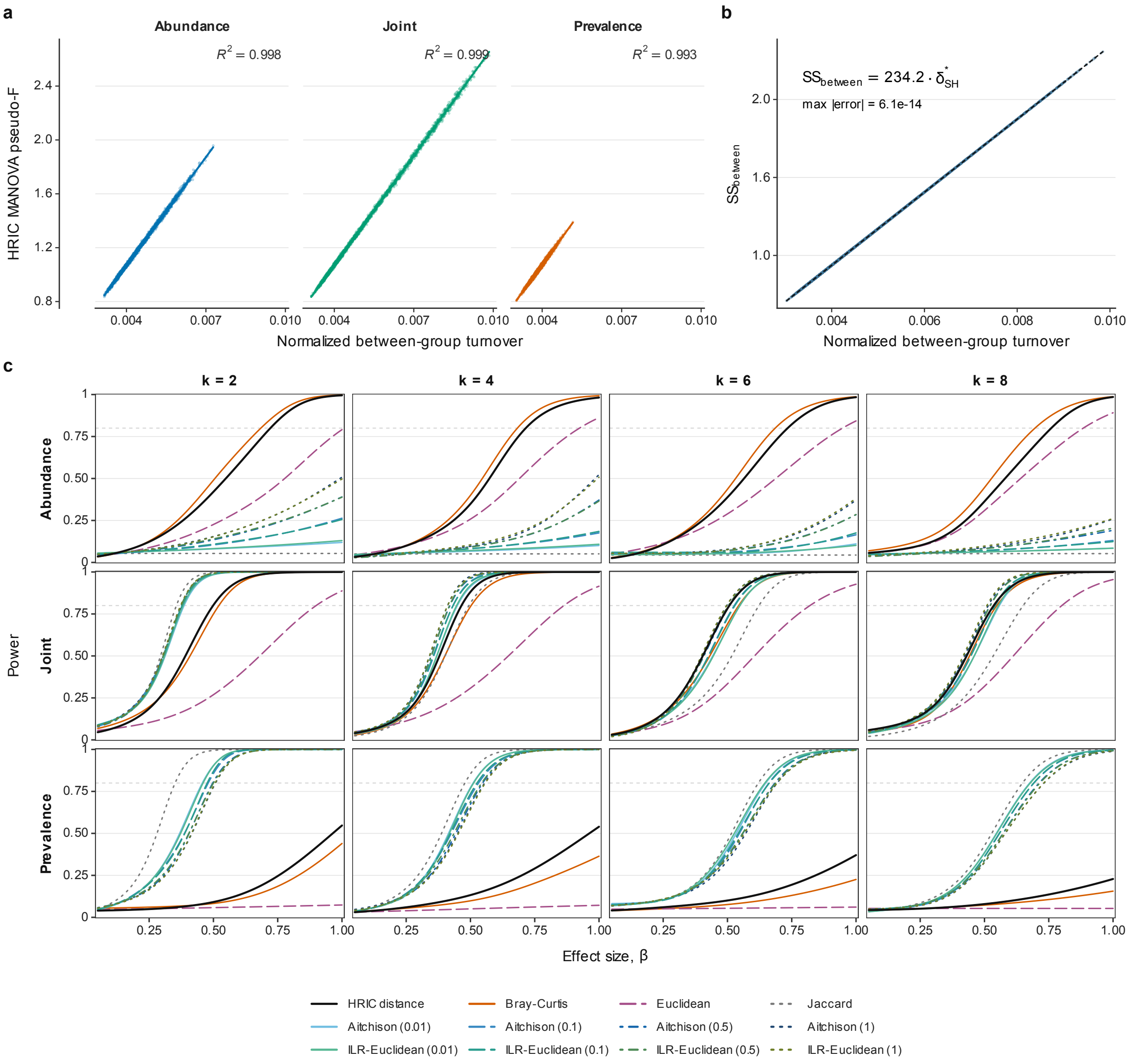

Supplementary Figure 1: **Condition-associated HRIC dispersion shares the between-condition geometry of PERMANOVA and shows signal-dependent power. a,** Relationship between normalized condition-associated HRIC dispersion, $\delta^*_{\mathrm{SH}} = \texttt{HRIC::SHdelta}(X, g)/A_p^2$, and the trace-based HRIC pseudo-$F$ statistic in the 12-cluster simulation. Each perturbation mode and effect size comprised 125 independent replicates, and quantities were averaged across the 12 possible perturbed clusters within each replicate. Points are replicate-level estimates; lines and shaded regions show mode-specific ordinary least-squares fits and 95% confidence intervals, with $R^2$ reported in each panel. **b,** Exact relationship between normalized condition-associated HRIC dispersion and $SS_{\text{between}}$. Points are replicate-level estimates, and the black dashed line is the theoretical identity. For $n = 100$ samples and $p = 616$ taxa, $nA_p^2 = 234.2$; the largest absolute numerical error in $SS_{\text{between}} = nA_p^2\delta^*_{\mathrm{SH}}$ was $6.1 \times 10^{-14}$. **c,** Power of PERMANOVA based on HRIC dissimilarity and comparator dissimilarities under abundance-only, joint abundance–prevalence and prevalence-only perturbations. Taxa were divided into $k = 2, 4, 6$ or 8 phylogenetically coherent clusters. For each $k$, the displayed perturbed cluster contained the largest number of taxa (308, 154, 103 and 77, respectively), with ties resolved by fitted total abundance. Each setting comprised 100 independently simulated datasets of 100 samples. Curves are binomial generalized additive model fits to empirical rejection proportions, and horizontal grey lines mark 80% power. HRIC dissimilarity is the Euclidean distance between HRIC coordinate vectors. Parenthetical values for Aitchison and ilr–Euclidean dissimilarities denote the pseudocount added before log-ratio transformation.

### Supplementary Note 4: Worked example and software implementation

#### S4.1 Worked diversity example

Consider three equally weighted communities on five species,

$$\begin{aligned}\boldsymbol{\pi}_A &= (1/3, 1/3, 1/3, 0, 0)^\mathsf{T},\\ \boldsymbol{\pi}_B &= (0, 1/3, 0, 1/3, 1/3)^\mathsf{T},\\ \boldsymbol{\pi}_C &= (1/3, 0, 1/3, 0, 1/3)^\mathsf{T}.\end{aligned} \tag{S71}$$

All three compositions lie on boundary faces of the five-part simplex, yet every HRIC coordinate is finite. Their order-2 Hill diversities are each 3. Pooling the relative abundances with equal community weights gives

$$\bar{\boldsymbol{\pi}} = (2/9, 2/9, 2/9, 1/9, 2/9)^\mathsf{T}, \tag{S72}$$

whose order-2 Hill diversity is 81/17. On the order-2 Hill-number scale, the additive excess is $81/17 - 3 = 30/17$, whereas multiplicative differentiation is $(81/17)/3 = 27/17$ [1, 3, 4]. The multiplicative value therefore corresponds to approximately 1.59 effectively distinct communities of effective local size 3.

The Simplex Hellinger quantities use the same three local compositions but a different, geometry-derived diversity scale. Their pooled regional composition is obtained from the normalized average of $\sqrt{\boldsymbol{\pi}_A}$, $\sqrt{\boldsymbol{\pi}_B}$ and $\sqrt{\boldsymbol{\pi}_C}$, as defined in Equation (S7), rather than by pooling raw reads or using the arithmetic composition in Equation (S72). Local alpha diversity, gamma diversity of the pooled regional composition and additive regional beta diversity then follow from Equations (S43), (S47) and (S48). The resulting regional beta component is non-negative by Theorem S3.5. The example also illustrates the fixed-dictionary requirement: all five species, including local zeros, define the common regional simplex.

#### S4.2 Computation from a community table

Let

$$X = (x_{ij}) \in \mathbb{R}^{n\times p}_{\geq 0}$$

have positive row totals $N_i = \sum_{j=1}^{p} x_{ij}$. For each row, the implementation performs the following operations:

1. close the abundance vector to $\boldsymbol{\pi}_i = \boldsymbol{x}_i/N_i$;
2. compute $\boldsymbol{y}_i = \sqrt{\boldsymbol{\pi}_i}$ and $\boldsymbol{y}_0 = p^{-1/2}\mathbf{1}$;
3. compute $c_i = \boldsymbol{y}_i^\mathsf{T}\boldsymbol{y}_0$, restrict floating-point departures of $c_i$ to $[0, 1]$, and set $s_i = \sqrt{\max(0, 1 - c_i^2)}$;
4. evaluate the radial factor as 1 when $s_i = 0$ and as $\arcsin(s_i)/s_i$ otherwise; and
5. return the HRIC row $\boldsymbol{z}_i$ as the radial factor multiplied by $\boldsymbol{y}_i - c_i\boldsymbol{y}_0$.

The complete transformation requires $O(np)$ arithmetic and $O(np)$ storage for the returned matrix. It uses no pseudocount, imputation or zero replacement. For extremely small $s_i$, the expansion

$$\frac{\arcsin(s_i)}{s_i} = 1 + \frac{s_i^2}{6} + \frac{3s_i^4}{40} + O(s_i^6) \tag{S73}$$

can be used, although direct evaluation is stable at ordinary double precision when the $s_i = 0$ limit is handled explicitly.

The software closes non-negative count or abundance rows internally. Each sample therefore enters the geometry through its relative composition rather than its library size. Library size can enter a downstream statistical model as a precision or sampling-depth covariate when scientifically required, but it does not alter the compositional definition. Rows with zero total abundance have no closure and must be removed or resolved before transformation. Comparisons on a shared HRIC scale also require the same feature dictionary and feature ordering.

##### S4.3 Correspondence with software functions

The mathematical quantities correspond to package functions as follows:

| Function | Returned quantity |
|---|---|
| `HRIC(X)` | Rows $\boldsymbol{z}_i = \mathrm{HRIC}(\boldsymbol{\pi}_i)$ in the centred $p$-coordinate representation. |
| `RHRIC(X, reference)` | Reference-based coordinates in Equation (S38). |
| `SHalpha(X)` | Vector $\alpha_{\mathrm{SH}}(\boldsymbol{\pi}_i)$ from Equation (S43). |
| `SHgamma(X)` | Scalar $\gamma_{\mathrm{SH}}(\boldsymbol{\Pi})$ from Equation (S47). |
| `SHbeta(X)` | Scalar $\beta_{\mathrm{SH}}(\boldsymbol{\Pi})$ from Equation (S48). |
| `SHdelta(X, group)` | Unnormalized condition-associated HRIC dispersion $\delta_{\mathrm{SH}}$ from Equation (S58). |

The implementation clips only floating-point departures from mathematically valid ranges: $c_i$ is restricted to $[0, 1]$, alpha and gamma diversity are restricted to $[0, 1]$, and theoretically non-negative regional beta and condition-associated dispersion values are truncated at zero only when round-off produces a small negative value. These operations are numerical safeguards and do not alter the population definitions.

#### Supplementary Note 5: Data processing and statistical implementation for the empirical analyses

##### S5.1 Data sources, preprocessing and computational implementation

The ocean analysis used the publicly available Tara Oceans prokaryote-enriched metagenomic 16S/18S rRNA miTAG abundance tables described by Salazar et al. [22]. Profiles for 180 samples are available through the study companion site and EBI BioStudies accession `S-BSST297`; matched sample and environmental metadata are available in Tables W1–W6 and through PANGAEA (doi:10.1594/PANGAEA.875582) [23]. Samples assigned in the metadata to the Arctic Ocean ($n = 38$) or North Atlantic Ocean ($n = 24$) were retained. Samples with zero total counts and features absent from every sample in a regional analysis were removed. The final region-specific feature dictionaries contained 10,122 and 11,163 observed features in the Arctic and North Atlantic, respectively. Taxonomic summaries used the deepest non-missing annotation from genus to kingdom.

The auto-FMT analysis used the processed 16S rRNA gene feature table and sample metadata from Qiita study `12277`, corresponding to the randomized trial reported by Taur et al. [24]. After removal of empty samples and globally absent features, the dataset contained 498 stools from 25 evaluable randomized patients, comprising 14 auto-FMT recipients and 11 controls, and 6,647 features. Trial metadata supplied treatment arm, allo-HSCT day, neutrophil-engraftment day, randomization day and the inverse-Simpson value reported by the original investigators. No rarefaction or pseudocount was applied in either empirical analysis. Non-negative count matrices were supplied directly to the HRIC package, which performs closure internally.

Analyses were performed in R 4.5.0 using `HRIC` 0.1.0, `phyloseq` 1.52.0, `lme4` 2.0.1, `sandwich` 3.1.1, `ggplot2` 4.0.3, `dplyr` 1.2.1, `tidyr` 1.3.2 and `patchwork` 1.3.2. Base-R `stats` functions were used for linear models, correlations, $t$-tests, Fisher's exact tests and multiplicity adjustment. Random seeds were fixed before resampling and permutation procedures.

##### S5.2 Ocean local–regional diversity analysis

Each water sample was treated as a local community, and all samples within an oceanographic region formed its regional ensemble. HRIC alpha and gamma diversity and additive regional beta diversity were calculated separately for the Arctic and North Atlantic with `HRIC::SHalpha`, `HRIC::SHgamma` and `HRIC::SHbeta`. Gamma diversity used the equal-community-weighted pooled regional composition defined in Equation (S7). The signed local–regional diversity gap for sample $i$ was

$$\tau_i = \gamma_{\mathrm{SH}}(\mathbf{\Pi}) - \alpha_{\mathrm{SH}}(\boldsymbol{\pi}_i),$$

whose regional mean equals additive regional beta diversity. Samples were ordered by increasing $\tau_i$ for the integrated figure.

Six environmental variables were examined: depth, temperature, oxygen, chlorophyll $a$, phosphate and nitrate. Within each region, associations between the signed local–regional diversity gap and an environmental variable were calculated from all finite sample–covariate pairs using two-sided Spearman rank correlation with `stats::cor.test`, `method="spearman"` and `exact=FALSE`. Ordinary least-squares lines and 95% confidence bands were included as visual summaries and were not used to calculate the Spearman statistics. The six correlations within each region were exploratory, and their reported $P$ values were not adjusted for multiplicity.

##### S5.3 Ocean taxon-resolved analysis

For the stacked compositional profiles, counts were aggregated to the deepest available taxonomic label and converted to relative abundance. The 19 taxa with the largest mean relative abundance within each region were displayed, and all remaining taxa were combined as Others. For each of the six environmental variables, the aggregated count matrix was transformed with `HRIC::HRIC`; a separate univariable linear model was then fitted for each of the 19 displayed taxon coordinates using `stats::lm`. The two-sided coefficient test assessed a zero slope, and Benjamini–Hochberg adjustment was applied across the 19 taxa within each region and environmental variable.

Nitrate was selected for the taxon-resolved panel because it was associated with the largest number of taxon coordinates at $P_{\mathrm{BH}} < 0.05$ among the six displayed environmental variables. This post hoc selection determined which taxon-level associations were displayed, and the corresponding results were interpreted as exploratory. Mapped ocean currents supplied geographical context for the North Atlantic sampling locations and did not enter the fitted statistical models.

##### S5.4 Auto-FMT time scales, personal references and outcomes

Clinical day 0 denoted allo-HSCT infusion. For randomized comparisons, event time was defined as stool-collection day minus randomization day. Event day 0 therefore denoted auto-FMT in the treatment arm and randomization in the control arm. Exact event-day-0 stools were excluded from the primary analysis because their collection order relative to auto-FMT was unavailable.

The earliest observed pre-HSCT stool, defined by clinical day $< 0$, served as each patient's personal compositional reference. The nearest stool within event days $-28$ to $-1$ defined the pre-index state. Pairwise HRIC dissimilarity between each stool and the patient's personal reference quantified individualized compositional restoration; smaller values indicated closer return to the personal pre-HSCT community. HRIC alpha diversity quantified within-stool diversity in the same coordinate representation.

#### S5.5 Longitudinal models and primary estimand

The primary longitudinal dataset comprised 98 stools contributed by 24 of the 25 evaluable patients and collected during event days 1–60. HRIC alpha diversity was modelled with a linear mixed-effects model containing treatment arm, separate linear event-time terms for days 1–14 and 15–60, both arm-by-time interactions, personal pre-HSCT alpha diversity, nearest pre-index alpha diversity, randomization day and a patient random intercept. The model was fitted by maximum likelihood. Continuous baseline variables were centred at their cohort means before fitting.

A corresponding patient random intercept in the HRIC-dissimilarity model had an estimated variance of zero. Pairwise HRIC dissimilarity from the personal reference was therefore modelled by ordinary least squares with treatment arm, the same piecewise-linear event-time terms and arm-by-time interactions, personal pre-HSCT alpha diversity, pre-index HRIC dissimilarity and randomization day. Uncertainty for this model was calculated with patient-clustered HC2 covariance.

The primary estimand was the adjusted auto-FMT-minus-control contrast averaged over event days 1–30. It was calculated by trapezoidal integration of the daily fixed-effect contrast curve rather than by selecting a single follow-up day. Uncertainty was estimated from 4,999 nonparametric bootstrap resamples of patients, sampled with replacement within randomized arm; the complete longitudinal record of each selected patient was retained. Point contrasts at days 14, 30 and 60 used the fitted model covariance.

#### S5.6 Sensitivity and corroborative recovery analyses

Sensitivity analyses replaced the earliest pre-HSCT alpha diversity with the mean of all available pre-HSCT values and repeated the primary models after including event-day-0 stools. Pre-index trajectory comparisons assessed whether the arms differed before the index event.

An event-time difference-in-differences analysis was used as a corroborative assessment of residual arm separation immediately before the index event. For each outcome, the adjusted between-arm contrast at event day $-1$ was subtracted from the fitted post-index contrast. The resulting estimand measures the change in arm separation relative to the common day-$-1$ anchor. Its interpretation requires a local parallel-trends condition: without auto-FMT, the adjusted arm difference would have remained stable over the short interval spanning day $-1$ and early follow-up. Randomization limits systematic treatment selection, while adjustment for personal pre-HSCT state, nearest pre-index state and randomization day accounts for measured differences in recovery stage. Because pre-index stools were sparse and irregularly timed, the difference-in-differences analysis corroborated the primary randomized-arm comparison rather than defining a separate primary estimand.

The 80% personal-baseline recovery analysis was descriptive. It was restricted to patients whose nearest pre-index HRIC alpha diversity was below 80% of their personal pre-HSCT value. Recovery required a threshold crossing by day 30 that was confirmed at the next observed sample, and arm-specific proportions were compared using a two-sided Fisher's exact test.

##### S5.7 Landmark diversity summaries and taxon contributions

One stool per patient was selected nearest the target day within each prespecified window: pre-index (days −28 to −1, target −1), early post-index (days 1–14, target 1) and later post-index (days 15–60, target 30). HRIC alpha and gamma diversity and additive regional beta diversity were calculated separately by randomized arm. Treatment-arm-associated HRIC dispersion at each landmark was calculated with `HRIC::SHdelta`; these landmark quantities were descriptive.

A taxon's contribution to alpha-diversity recovery was defined as its pre-index contribution to $1 - \alpha$ minus its contribution at the corresponding post-index landmark, averaged within arm. Because the within-sample contribution of taxon $j$ is $z_{ij}^2 / A_p^2$, positive recovery values indicate that the taxon contributed less to community unevenness at follow-up. Contributions were aggregated to the deepest available taxonomic annotation.

##### S5.8 Comparison with inverse-Simpson diversity and Pielou evenness

The alpha-diversity index reported by Taur et al. [24] was recomputed as

$$\log \left( \frac{1}{\sum_{j=1}^{p} \pi_{ij}^2} \right), \tag{S74}$$

where $\pi_{ij}$ is the relative abundance of taxon $j$ in sample $i$. Pielou evenness was calculated as

$$\frac{-\sum_{j=1}^{p} \pi_{ij} \log(\pi_{ij})}{\log(S_i)}, \tag{S75}$$

where $S_i$ is observed richness. Concordance with HRIC alpha diversity was summarized by Spearman correlation across all 498 stools.

Consecutive stools were ordered by clinical day within patient, and adjacent transitions with opposite directions under HRIC alpha and either conventional index were decomposed exactly. The HRIC contribution of taxon $j$ to a transition from a before sample to an after sample was

$$\frac{z_{ij,\text{before}}^2 - z_{ij,\text{after}}^2}{A_p^2}. \tag{S76}$$

Changes in log inverse-Simpson diversity were localized through the corresponding changes in $\pi_{ij}^2$. Changes in Pielou evenness were partitioned into taxon-specific entropy terms and a separate term for the change in the $\log(S_i)$ denominator. Taxon contributions were aggregated to the deepest available annotation.

##### S5.9 Supplementary auto-FMT results

Pre-index trajectory comparisons showed limited evidence of differential recovery before the index event. The between-arm difference in the pre-index alpha-diversity slope had $P = 0.058$, and the corresponding comparison for pairwise HRIC dissimilarity from the personal pre-HSCT reference had $P = 0.373$. In the corroborative event-time difference-in-differences analysis, the early-window auto-FMT-minus-control contrast relative to the adjusted day-−1 arm difference was +0.0580 for HRIC alpha diversity (95% model-based confidence interval, +0.0350 to +0.0809) and −0.603 for pairwise HRIC dissimilarity from the personal reference (95% model-based confidence interval, −0.858 to −0.348). In the descriptive threshold analysis, 6 of 13 auto-FMT recipients and 1 of 8 controls whose nearest pre-index alpha diversity was below 80% of their personal pre-HSCT value had a crossing confirmed at the next observed sample by day 30 (two-sided Fisher's exact $P = 0.174$).

Across all 498 stools, HRIC alpha diversity was strongly concordant with log inverse-Simpson diversity (Spearman's $\rho = 0.913$) and Pielou evenness ($\rho = 0.862$). Eight adjacent-sample transitions in patients 0020 and 0061 nevertheless changed in opposite directions under HRIC alpha and at least one conventional index. In patient 0020 between clinical days 5 and 12, [Coprobacillaceae] decreased from 77.4% to undetectable, *Lactobacillus* increased from 11.0% to 80.7%, and observed richness decreased from 39 to 15. Log inverse-Simpson diversity and Pielou evenness increased by 0.593 and 0.222, respectively, whereas HRIC alpha diversity decreased by 0.0019. Across the discordant transitions, recurrent contributors included *Lactobacillus*, [Coprobacillaceae], *Streptococcus*, *Blautia*, [*Eubacterium*], Ruminococcaceae, *Bifidobacterium* and *Akkermansia*. In patient 0061, several reversals involving Pielou evenness arose principally from changes in its log-richness denominator. These localized disagreements were therefore interpretable consequences of how the indices weight dominant taxa, the low-abundance tail and richness normalization.

The largest early reductions in taxon contributions to $1 - \alpha$ in the auto-FMT arm involved Listeriaceae and Enterobacteriaceae, followed by smaller reductions for *Lactococcus*, *Streptococcus* and *Lactobacillus*. Across the union of the 50 taxa with the largest absolute contributions in either arm, the arm-specific profiles had Spearman correlations of 0.58 in the early window and 0.49 in the later window. Five taxa were shared between the arm-specific top-10 sets at each landmark. These summaries support a partly shared but substantially patient- and arm-specific pattern of taxonomic contribution to alpha-diversity recovery.

### References

[1] Robert H. Whittaker. Evolution and measurement of species diversity. *Taxon*, 21(2–3):213–251, 1972. doi: 10.2307/1218190.

[2] Russell Lande. Statistics and partitioning of species diversity, and similarity among multiple communities. *Oikos*, 76(1):5–13, 1996. doi: 10.2307/3545743.

[3] Mark O. Hill. Diversity and evenness: A unifying notation and its consequences. *Ecology*, 54(2): 427–432, 1973. doi: 10.2307/1934352.

[4] Lou Jost. Partitioning diversity into independent alpha and beta components. *Ecology*, 88(10): 2427–2439, 2007. doi: 10.1890/06-1736.1.

[5] J. Roger Bray and John T. Curtis. An ordination of the upland forest communities of southern wisconsin. *Ecological Monographs*, 27(4):325–349, 1957. doi: 10.2307/1942268.

[6] Hugh G. Gauch, Jr. and Robert H. Whittaker. Comparison of ordination techniques. *Ecology*, 53(5):868–875, 1972. doi: 10.2307/1934302.

[7] Patricia Koleff, Kevin J. Gaston, and Jack J. Lennon. Measuring beta diversity for presence–absence data. *Journal of Animal Ecology*, 72(3):367–382, 2003. doi: 10.1046/j.1365-2656.2003.00710.x.

[8] Hanna Tuomisto. A diversity of beta diversities: straightening up a concept gone awry. part 1. defining beta diversity as a function of alpha and gamma diversity. *Ecography*, 33(1):2–22, 2010. doi: 10.1111/j.1600-0587.2009.05880.x.

[9] Pierre Legendre, Daniel Borcard, and Pedro R. Peres-Neto. Analyzing beta diversity: Partitioning the spatial variation of community composition data. *Ecological Monographs*, 75(4):435–450, 2005. doi: 10.1890/05-0549.

[10] Marti J. Anderson, Kari E. Ellingsen, and Brian H. McArdle. Multivariate dispersion as a measure of beta diversity. *Ecology Letters*, 9(6):683–693, 2006. doi: 10.1111/j.1461-0248.2006.00926.x.

[11] Carlo Ricotta and Sabina Burrascano. Testing for differences in beta diversity with asymmetric dissimilarities. *Ecological Indicators*, 9(4):719–724, 2009. doi: 10.1016/j.ecolind.2008.09.003.

[12] Pierre Legendre and Miquel De Cáceres. Beta diversity as the variance of community data: Dissimilarity coefficients and partitioning. *Ecology Letters*, 16(8):951–963, 2013. doi: 10.1111/ele.12141.

[13] Brian H. McArdle and Marti J. Anderson. Fitting multivariate models to community data: A comment on distance-based redundancy analysis. *Ecology*, 82(1):290–297, 2001. doi: 10.1890/0012-9658(2001)082[0290:FMMTCD]2.0.CO;2.

[14] Marti J. Anderson. A new method for non-parametric multivariate analysis of variance. *Austral Ecology*, 26(1):32–46, 2001. doi: 10.1111/j.1442-9993.2001.01070.pp.x.

[15] Ni Zhao, Jun Chen, Ian M. Carroll, Tamar Ringel-Kulka, Michael P. Epstein, Hua Zhou, Jin J. Zhou, Yehuda Ringel, Hongzhe Li, and Michael C. Wu. Testing in microbiome-profiling studies with MiRKAT, the microbiome regression-based kernel association test. *The American Journal of Human Genetics*, 96(5):797–807, 2015. doi: 10.1016/j.ajhg.2015.04.003.

[16] Calyampudi Radhakrishna Rao. Information and the accuracy attainable in the estimation of statistical parameters. *Bulletin of the Calcutta Mathematical Society*, 37(3):81–91, 1945.

[17] John Aitchison. The statistical analysis of compositional data. *Journal of the Royal Statistical Society: Series B (Methodological)*, 44(2):139–177, 1982. doi: 10.1111/j.2517-6161.1982.tb01195.x.

[18] Juan José Egozcue, Vera Pawlowsky-Glahn, Glòria Mateu-Figueras, and Carles Barceló-Vidal. Isometric logratio transformations for compositional data analysis. *Mathematical Geology*, 35(3):279–300, 2003. doi: 10.1023/A:1023818214614.

[19] Emily S. Charlson, Jun Chen, Rebecca Custers-Allen, Kyle Bittinger, Hongzhe Li, Rohini Sinha, Jennifer Hwang, Frederic D. Bushman, and Ronald G. Collman. Disordered microbial communities in the upper respiratory tract of cigarette smokers. *PLoS ONE*, 5(12):e15216, 2010. doi: 10.1371/journal.pone.0015216.

[20] Jun Chen, Kyle Bittinger, Emily S. Charlson, Christian Hoffmann, James Lewis, Gary D. Wu, Ronald G. Collman, Frederic D. Bushman, and Hongzhe Li. Associating microbiome composition with environmental covariates using generalized UniFrac distances. *Bioinformatics*, 28(16):2106–2113, 2012. doi: 10.1093/bioinformatics/bts342.

[21] Siyuan Ma, Boyu Ren, Himel Mallick, Yo Sup Moon, Emma Schwager, Sagun Maharjan, Timothy L. Tickle, Yiren Lu, Rachel N. Carmody, Eric A. Franzosa, Lucas Janson, and Curtis Huttenhower. A statistical model for describing and simulating microbial community profiles. *PLoS Computational Biology*, 17(9):e1008913, 2021. doi: 10.1371/journal.pcbi.1008913.

[22] Guillem Salazar et al. Gene expression changes and community turnover differentially shape the global ocean metatranscriptome. *Cell*, 179(5):1068–1083.e21, 2019. doi: 10.1016/j.cell.2019.10.014.

[23] Stéphane Pesant et al. Open science resources for the discovery and analysis of Tara Oceans data. *Scientific Data*, 2:150023, 2015. doi: 10.1038/sdata.2015.23.

[24] Ying Taur, Katharine Coyte, Jonas Schluter, Elizabeth Robilotti, Clara Figueroa, Mergim Gjonbalaj, Eric R. Littmann, Lilan Ling, Liana Miller, Yonten Gyaltshen, Emily Fontana, Sejal Morjaria, Boglarka Gyurkocza, Miguel-Angel Perales, Hugo Castro-Malaspina, Roni Tamari, Doris Ponce, Guenther Koehne, Juliet Barker, Ann Jakubowski, Esperanza Papadopoulos, Parastoo Dahi, Craig Sauter, Brian Shaffer, James W. Young, Jonathan U. Peled, Regina C. Meagher, Robert R. Jenq, Marcel R. M. van den Brink, Sergio A. Giralt, Eric G. Pamer, and Joao B. Xavier. Reconstitution of the gut microbiota of antibiotic-treated patients by autologous fecal microbiota transplant. *Science Translational Medicine*, 10(460):eaap9489, 2018. doi: 10.1126/scitranslmed.aap9489.
